# Large, stable spikes exhibit differential broadening in excitatory and inhibitory neocortical boutons

**DOI:** 10.1101/2020.10.19.346346

**Authors:** Andreas Ritzau-Jost, Timur Tsintsadze, Martin Krueger, Jonas Ader, Ingo Bechmann, Jens Eilers, Boris Barbour, Stephen M. Smith, Stefan Hallermann

## Abstract

Presynaptic action potential spikes control neurotransmitter release and thus interneuronal communication. However, the properties and the dynamics of presynaptic spikes in the neocortex remain enigmatic because boutons in the neocortex are small and direct patch-clamp recordings have not been performed. Here we report direct recordings from boutons of neocortical pyramidal neurons and interneurons. Our data reveal rapid and large presynaptic action potentials in layer 5 neurons and fast-spiking interneurons reliably propagating into axon collaterals. For in-depth analyses we validate boutons of mature cultured neurons as models for excitatory neocortical boutons, demonstrating that the presynaptic spike amplitude was unaffected by potassium channels, homeostatic long-term plasticity, and high-frequency firing. In contrast to the stable amplitude, presynaptic spikes profoundly broadened for example during high-frequency firing in layer 5 pyramidal neurons but not in fast-spiking interneurons. Thus, our data demonstrate large presynaptic spikes and fundamental differences between excitatory and inhibitory boutons in the neocortex.

## INTRODUCTION

Spikes propagating along the axon trigger neurotransmitter release at presynaptic boutons (Bean, 2007). The amplitude and the duration of the presynaptic spike critically control the gating of voltage-dependent calcium channels (Borst and Sakmann, 1998; Sabatini and Regehr, 1997) and thereby the strength of synaptic transmission (Sakaba and Neher, 2001; Wheeler et al., 1994; Zbili et al., 2020). Therefore, alterations in the shape of the presynaptic action potential contribute to short- and long-term plasticity in the hippocampus (Carta et al., 2014; Geiger and Jonas, 2000). However, fundamental properties and the plasticity of presynaptic spikes in the neocortex remain unclear because boutons in the neocortex are small (diameter ~0.7 μm; Rollenhagen et al., 2018) and thus difficult to investigate directly. To overcome these limitations in the understanding of fundamental properties of the central nervous system (CNS), we established direct presynaptic recordings from two prototypical neurons in the neocortex. We chose layer 5 pyramidal neurons because they are among the best-studied neurons of the CNS and direct recordings from their dendrites (Nevian et al., 2007; Stuart and Sakmann, 1994) and main axon (Cohen et al., 2020; Shu et al., 2006) have contributed substantially to our understanding of neuronal function in general (reviewed in Kole and Stuart, 2012; Stuart et al., 2016). In comparison, we focused on the presynaptic spike of fast-spiking neocortical interneurons, one of the main sources of GABAergic inhibition in the neocortex (Markram et al., 2004; Tremblay et al., 2016).

Recordings of presynaptic action potentials from large boutons in the brainstem (Forsythe, 1994; Sierksma and Borst, 2017), hippocampus (Alle et al., 2009; Geiger and Jonas, 2000), pituitary gland (Jackson et al., 1991), and cerebellum (Ritzau-Jost et al., 2014) revealed large and stable action potentials, although deviations from an “all-or-none” behavior occur due to changes in the steady-state sodium and potassium channel availability (Alle and Geiger, 2006; Johnston et al., 2009; reviewed in Zbili and Debanne, 2019). However, recordings from small conventional boutons, forming >99% of all central synapses, were often restricted to the cell-attached configuration preventing the measurement of the presynaptic action potential amplitude (Rowan et al., 2014; Sasaki et al., 2011; Smith et al., 2004). Whole-cell recordings have only been possible at small inhibitory boutons in acute brain slices of the cerebellum (Begum et al., 2016; Kawaguchi and Sakaba, 2015; Southan and Robertson, 1998) and small boutons of cultured hippocampal neurons (Vivekananda et al., 2017), but AP amplitude was not quantified. Whole-cell recordings in axon shafts and cut axons (blebs) of inhibitory interneurons in acute brain slices of the hippocampus revealed large action potentials (Hu et al., 2018). In contrast, attempts to quantify the amplitude of action potentials based on recordings with voltage sensitive dyes in boutons of cultured hippocampal neurons suggested small amplitudes (~70% of the somatic action potential amplitude) that were dynamically regulated during plasticity (Hoppa et al., 2014).

Here, by performing direct recordings from small boutons of neocortical neurons, we report that the amplitude of the presynaptic action potential is large and surprisingly reliable with no evidence of conduction block during high-frequency firing. In contrast to the constancy of the spike amplitude, the presynaptic spike duration changed during various interventions, including broadening during short-term activity in excitatory but not inhibitory boutons.

## RESULTS

### Brief action potentials in boutons of layer 5 pyramidal neurons

To visualize the axonal arbor and *en passant* presynaptic boutons of layer 5 pyramidal neurons in acute brain slices, somatic whole-cell recordings with fluorescent pipette solution were obtained (Figure 1A). Somatic action potentials were elicited and action potential-evoked currents were simultaneously recorded in boutons with recording pipettes placed on their surface in loose-seal configuration (Figure 1B). Somatic action potentials showed half-durations of 656 ± 35 μs (measured from resting membrane potential, n = 32; Figure 1C). In loose-seal bouton-recordings, peak-to-peak duration of biphasic action potential-evoked currents (379 ± 25 μs) was substantially shorter than somatic half-durations (P < 0.001; n = 32; Figure 1C). As a first approximation, the extracellular current reflects the first derivative of the membrane voltage (Bean, 2007) and thus the peak-to-peak duration is a measure of the spike half-duration. The ohmic current components and the extracellular volley of the propagating action potential (Sabatini and Regehr, 1997) could also contribute to our loose-seal signal. However, comparison of loose-seal and whole-cell recordings from both the soma and the bouton showed that the peak-to-peak duration of loose-seal recorded currents reflected the action potential half-duration (Figures S1A – C; see below, Figure S4C). Similar to boutons of principal neurons, we evaluated boutons of neocortical fast-spiking interneurons (Figure 1D) that are known to have short action potentials (Casale et al., 2015). Peak-to-peak durations in interneuron boutons were shorter compared to those in pyramidal neurons (Figures 1E, and F) and similar to somatic action-potential duration (consistent with voltage-sensitive dye recordings; Casale et al., 2015). Thus, loose-seal recordings indicate that boutons of layer 5 pyramidal neurons have shorter action potentials compared to the soma.

**Figure 1.**
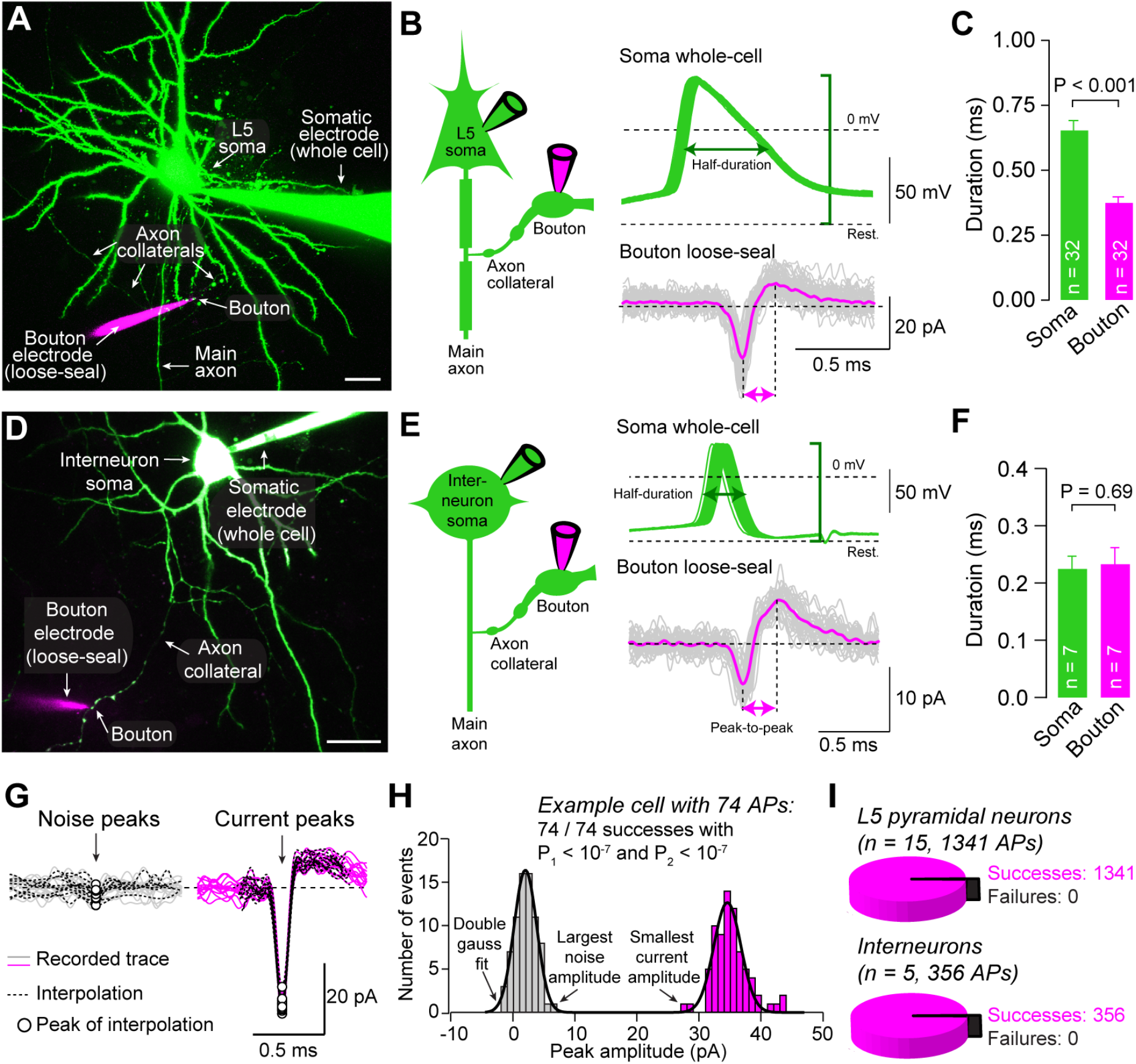
Brief action potentials in boutons of layer 5 pyramidal neurons. **(A)** Two-photon-(2P-)image of a neocortical layer 5 pyramidal neuron recorded in wholecell mode (green; 20 μM Atto-488 in the somatic pipette; maximum z-projection of a stack of 106 images, z-step size 1.5 μm). A second pipette (magenta; 200 μM Atto-594 in pipette) simultaneously recorded action potential-evoked currents from a bouton in loose-seal mode. Scale bar 20 μm. **(B)** *Left:* Pictogram of the recording configuration. *Right:* Overlay of 50 evoked somatic action potentials (green) and currents in a simultaneous loose-seal bouton recording (single traces in grey, average in magenta). Somatic action potential amplitude (green bracket), somatic action potential half-duration (green arrow), and duration of loose-seal recorded currents from in bouton recordings (magenta arrows) indicated. **(C)** Half-duration of somatic action potentials and peak-to-peak current duration in boutons during paired bouton-soma recordings from pyramidal neurons (bar graphs as mean ± SEM, color code as in B, n indicates number of paired soma-bouton recordings). **(D)** 2P-image of a paired bouton-soma recording from a neocortical interneuron (maximum z-projection of a stack of 57 images, z-step size 1.0 μm). Scale bar 20 μm. **(E)** *Left:* Pictogram of the recording configuration. *Right:* overlay of 32 evoked somatic action potentials (green) and simultaneously recorded currents in loose-seal mode from a bouton (single traces in grey, average in magenta). **(F)** Half-duration of somatic action potentials and peak-to-peak current duration in boutons during paired bouton-soma recordings from interneurons (bar graphs as mean ± SEM, n indicates number of paired soma-bouton recordings). **(G)** Overlay of 15 exemplary action potential-evoked currents from a loose-seal bouton recording of a pyramidal neuron. Noise and sodium current peak amplitudes (dots) are determined by interpolations (dashed lines). **(H)** Histograms of sodium and noise peak amplitudes fitted with double-gaussian fits. From the largest noise peak and the smallest current peak, two probabilities related to the reliability of action potential detection (*P*_1_ and *P*_2_) were calculated (see *methods*). **(I)** Proportion of failures of action potential propagation in paired soma-bouton recordings from pyramidal neurons and interneurons (n = 15 and 5, respectively). All somatically evoked action potentials (n = 1341 for pyramidal neurons, n = 356 for interneurons) were detected in loose-seal bouton recordings and no action potential propagation failure occurred. See also Figure S1.

Bouton recording sites were ~100 to 300 μm away from the offset of the main axon from the soma and separated from the main axon by 1 to 6 branch points (n = 19; Figures S1D and E). Peak-to-peak durations recorded in loose-seal mode were independent from distance of recording sites and branch point number (correlation coefficients 0.067 and −0.055, respectively; Figures S1D and E). The geometry of axonal branch points reduces the safety factor for action potential propagation marking them out as potential sites for conduction failure (Swadlow et al., 1980). However, we found that somatically elicited action potentials always propagated beyond branch points in recordings from layer 5 pyramidal neurons (15 bouton-soma pairs; total of 1341 action potentials; Figures 1G – I), in agreement with previous reports of reliable spike invasion into collaterals of neocortical pyramidal neurons and cultured neocortical neurons (Cox et al., 2000; Hamada et al., 2017; Popovic et al., 2011; Radivojevic et al., 2017). Similarly, spikes successfully propagated into collaterals of neocortical interneurons (six bouton-soma pairs; total of 356 action potentials; Figure 1I). We confirmed that these were not false positives by comparing the peak noise in the presence and absence of evoked action potentials (Figure 1G). From the complete absence of overlap in the distributions of peak events (Figure 1H) we estimated the probability of false-positively detecting an action potential to be at least below 10^-4^ in all experiments (see *methods*). This indicates that propagation of single action potentials into axon collaterals of layer 5 pyramidal neurons and interneurons is reliable and unimpeded by branch points.

### Discrepancy between whole-cell and loose-seal bouton recordings

Loose-seal recordings reliably reported action potential time-course, but did not provide information about action potential amplitude. To permit direct whole-bouton recordings from layer 5 pyramidal neurons we first compared patch pipettes fabricated from borosilicate and quartz glass. Quartz pipettes have favorable electrical characteristics (Benndorf, 1995; Dudel et al., 2000) and geometrical properties likely to facilitate recordings from small structures (Figure 2A and Figure S2). Therefore, quartz pipettes were used for recordings from boutons. Boutons were visualized by prior somatic filling with green fluorescent solution and successful whole-bouton configuration was confirmed by the diffusion of red fluorophore from the bouton pipette into the green-labeled axon (Figure 2B) as well as functionally by presynaptic action potentials elicited by somatic action potentials (Figure 2C). Curiously, bouton action potentials reached lower peak potentials (0.0 ± 5.5 mV, n = 5; Figure 2D) and were broader (917 ± 87 μs, n = 5; Figure 2D) compared to somatic action potentials (P < 0.001 and P < 0.001, respectively). Surprisingly, action potentials recorded from boutons in whole-cell mode were also broader than those recorded in loose-seal mode (P < 0.001; cf. Figure 1C). Consistently, action potentials in boutons of fast-spiking interneurons were broadened in whole-cell recordings compared to somatic action potentials (P < 0.03; Figures 2E – G) and compared to the loose-seal recordings (P < 0.03; cf. Figure 1F). Thus, in both, boutons of pyramidal neurons and interneurons, action potentials were broader in whole-cell recordings compared with loose-seal recordings. It is well established that compensation of the pipette capacitance critically impacts action potentials in whole-cell recordings (Barbor, 2014; Brette and Destexhe, 2012; Purves, 1981). We therefore hypothesized that whole-cell recordings interfered with action potentials in these small boutons.

**Figure 2.**
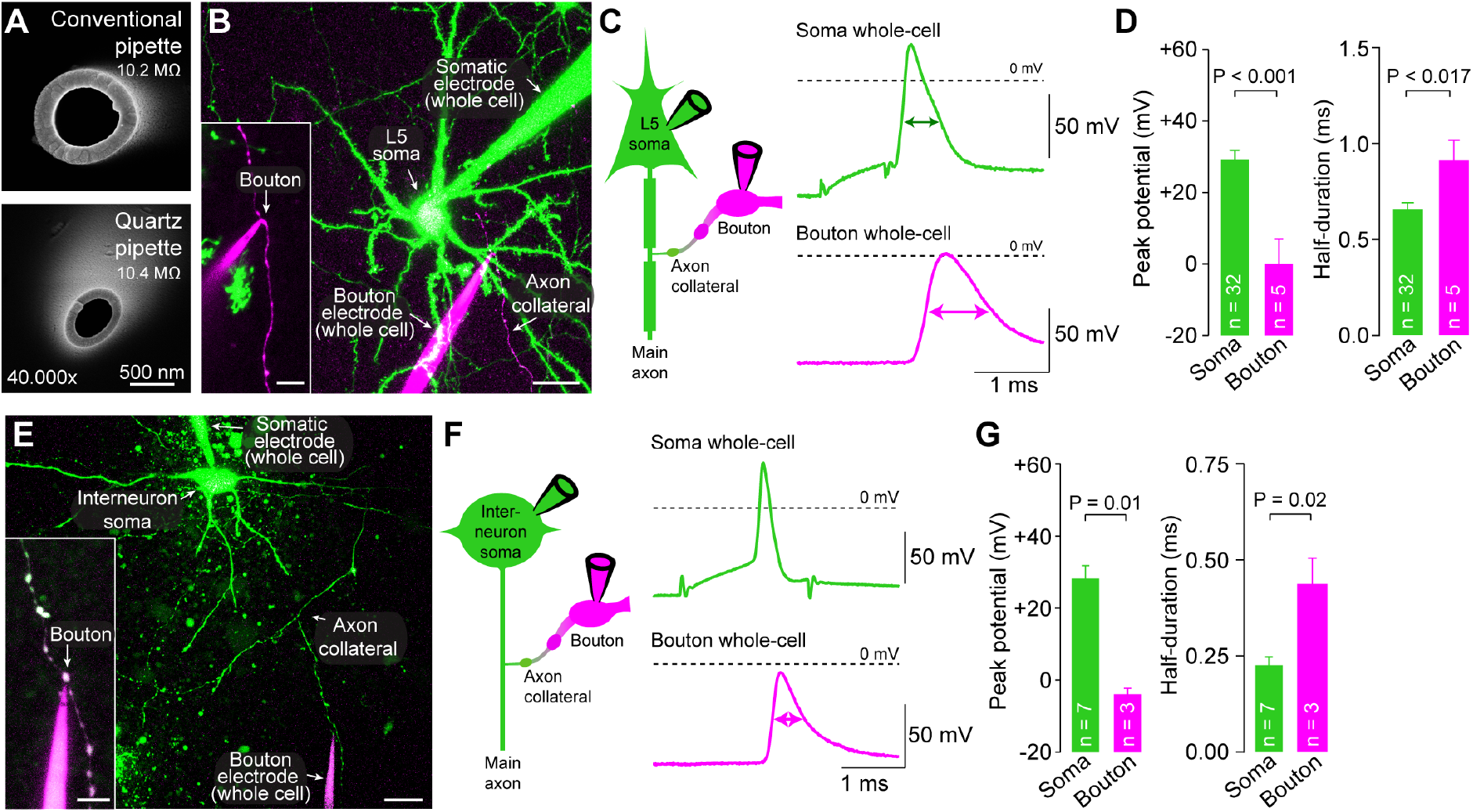
Discrepancy between whole-cell and loose-seal bouton recordings. **(A)** *Top:* Electron microscopic image of the tip of a borosilicate glass pipette. *Bottom:* Electron microscopic image of a quartz glass pipette of similar pipette resistances. **(B)** 2P-image of a paired whole-cell recording from the soma (green pipette) and bouton (magenta pipette) of a layer 5 pyramidal neuron. Scale bar 20 μm, maximum z-projection of a stack of 57 images, z-step size 1.25 μm. *Inset:* Magnified whole-cell bouton recording, scale bar 5 μm, maximum z-projection of a stack of 23 images, z-step size 0.5 μm. **(C)** *Left:* Pictogram of the recording configuration. *Right:* Example whole-cell recorded somatic (green) and bouton action potentials (magenta) upon somatic current-injection. Action potential half-durations in somatic and bouton recordings are indicated by arrows. **(D)** *Left:* Peak potential of somatic and bouton action potentials in paired bouton-soma recordings form layer 5 pyramidal neurons. *Right:* Half-duration of somatic and bouton action potentials in paired recordings form layer 5 pyramidal neurons (bar graphs as mean ± SEM, color code as in B, n indicates number of individual paired soma-bouton recordings). **(E)** 2P-image of a paired whole-cell recording from the soma (green pipette) and bouton (magenta pipette) of a neocortical interneuron. Scale bar 20 μm, maximum z-projection of a stack of 92 images, z-step size 1.0 μm. *Inset:* Magnified whole-cell bouton recording, scale bar 5 μm, maximum z-projection of a stack of 16 images, z-step size 0.5 μm. **(F)** *Left:* Pictogram of the recording configuration. *Right:* Whole-cell recorded somatic (green) and bouton action potentials (magenta) evoked by somatic current-injection. **(G)** Left: Peak potential of somatic and bouton action potentials in paired bouton-soma recordings form neocortical interneurons. *Right:* Half-duration of somatic and bouton action potentials in paired recordings form interneurons (bar graphs as mean ± SEM, color code as in F). See also Figure S2.

### Pipette capacitance critically impacts whole-bouton action potential measurements

To investigate potential errors in whole-cell recordings at small structures, we developed an electrical circuit with a capacitance estimated to be equivalent to that of an *en passant* bouton (capacitance of 0.5 pF) and a typical patch pipette including amplifier head-stage and the pipette holder (capacitance of 7.5 pF; Figure 3A; for passive properties of boutons also see Figure S3A). Rectangular command voltages (V_in_) were transformed into artificial action potential waveforms (‘target’) by the RC characteristics of the bouton-equivalent circuit and measured by an on-board operational amplifier (in the following referred to as V_m_; Fig 3A, B). Connecting the pipette equivalent circuit with the attached current-clamp amplifier distorted both the action potential waveforms recorded by the current-clamp amplifier (V_cc_) and V_m_ (Figure 3B). V_cc_ and V_m_ were undistorted only when the capacitance neutralization circuit of the amplifier could fully compensate the pipette capacitance (Figure 3B).

**Figure 3.**
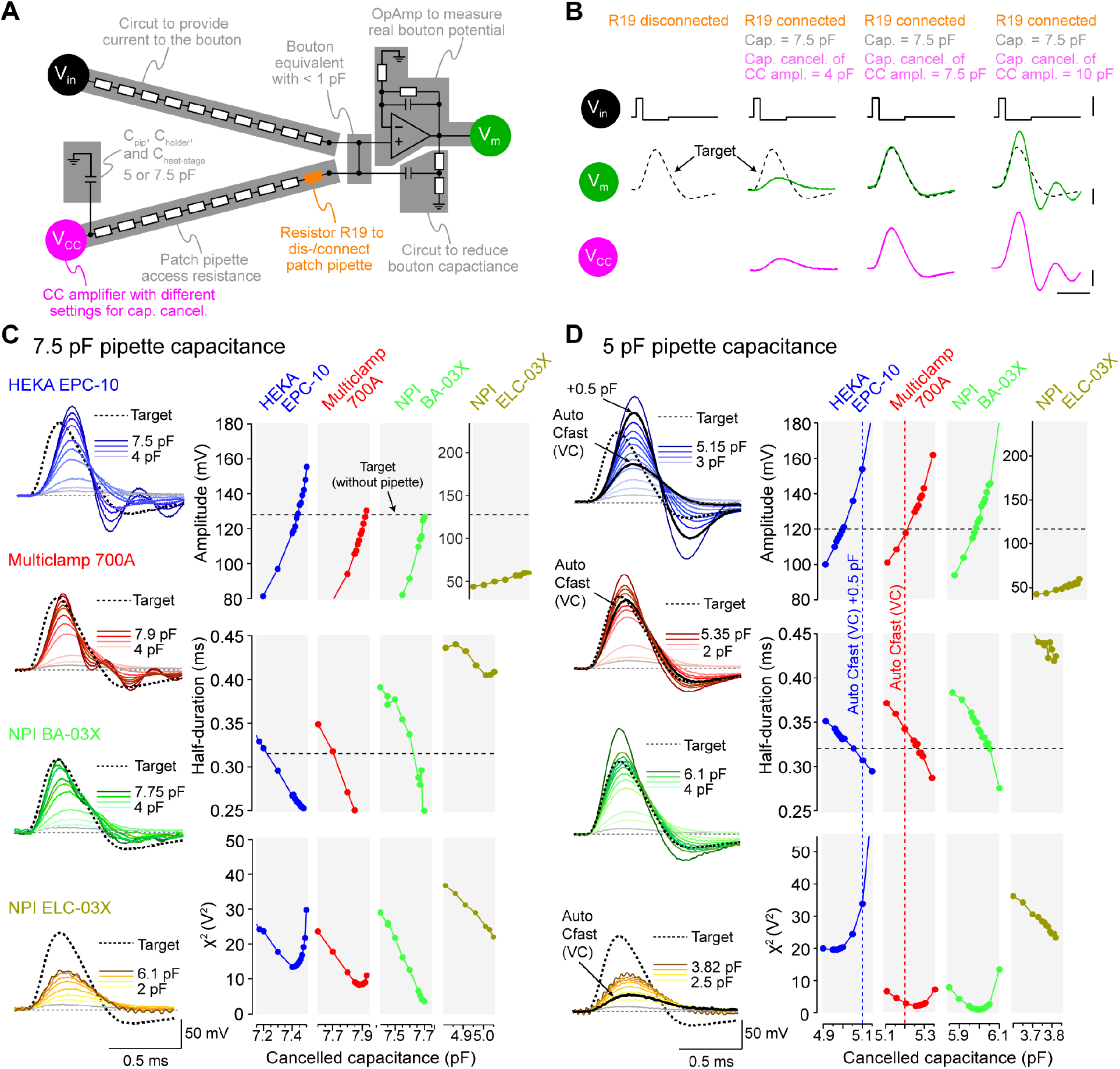
Pipette capacitance critically impacts whole-bouton action potential measurements. **(A)** Illustration of the test circuit used to determine the effect of pipette capacitance and pipette capacitance cancellation on action potential recordings. Command voltages are provided to the circuit at V_in_ and transformed by the circuit’s RC characteristics into action potential-like responses. These are recorded either directly (V_m_) after amplification by an on-board operational amplifier (OpAmp; ~0.5 pF input capacitance) or by a current-clamp amplifier (V_cc_) connected via an electronic circuit simulating the patch pipette’s passive electrical properties. **(B)** Illustration of the test procedure applied to determine the impact of pipette capacitance cancellation on presynaptic action potentials. Voltage commands are first applied to the circuit (at V_in_) when the pipette capacitance and current-clamp amplifier are disconnected (R19 disconnected) to evoke action potential-like voltages at V_m_ (‘target’). After reconnection of the pipette capacitance and the current-clamp amplifier (R19 connected) action potentials are recorded at different pipette capacitance cancellation settings of the connected current-clamp amplifier. **(C)** *Left:* Voltages recorded with different current-clamp amplifiers (see color code) with different degrees of pipette capacitance cancellation applied to a ~7.5 pF pipette capacitance (compensated pipette capacitance coded by brightness of individual traces). *Right:* Amplitude, half-duration and sum of squared errors (*χ*^2^) of recorded action potentials over cancelled capacitance for individual current-clamp amplifiers. Amplitude and half-duration of the ‘target’ waveform indicated by horizontal dashed lines. **(D)** Recordings as in C but for ~5 pF pipette capacitance. Additionally, pipette capacitance values of the disconnected pipette branch (R19 disconnected in panel A) were determined with each individual amplifier that featured voltage-clamp mode by automated voltage-clamp routines and are indicated by vertical dashed lines. See also Figure S3.

To determine the size of errors we compared action potential amplitude and half-duration of recorded action potentials to the ‘target’ waveform (horizontal dashed lines, Figure 3C) following variable capacitance cancellation by four different amplifiers. Deviations between recorded and ‘target’ waveform were further quantified by their sum of squared errors (*χ*^2^). The setting of the pipette capacitance cancellation leading to optimal amplitude, half-duration, and did not match for most amplifiers and substantial differences in the performance of different amplifiers were observed (Figure 3C).

We explored if lowering the pipette capacitance (including the amplifier head-stage and the pipette holder) to 5 pF increased the accuracy of current-clamp recordings. Measurements performed with lower capacitance generally improved the coincidence of the optimal amplitude, half-duration, and *χ*^2^ measurements (Fig 3D). The Multiclamp 700A (a combined voltage- and current-clamp amplifier) and the NPI BA-03X (current-clamp only amplifier) were better able than other amplifiers to accurately measure the action potential amplitude and half-duration and reached minimal deviation due to voltage oscillation (Magistretti et al., 1996; Fig 3D). However, only a small range of capacitance cancellations provided correct amplitude and half-duration as well as minimal *χ*^2^, emphasizing the importance of precise capacitance cancellation for whole-cell recordings from small boutons.

Classically, in somatic current-clamp recordings capacitance cancellation is optimized by identifying the degree of cancellation at which voltage oscillations occur upon current injection (Marty and Neher, 2009; Figure S3B). However, for the relatively high pipette capacitance/cell capacitance ratio present in our *en passant* bouton model (~10/1) there was no clear onset of oscillations and small cancellation errors resulted in substantial distortion of the voltage deflection (Figure S3C), rendering this approach inadequate for correct pipette capacitance compensation. Instead, we used patch-clamp amplifiers that have been designed to permit both voltage-clamp and current-clamp recordings and employed the automated capacitance cancellation in voltage-clamp mode to determine the pipette capacitance in the cell-attached configuration (see *methods* section for details). Indeed, the value obtained in voltage-clamp mode (vertical dashed lines in Figure 3D) by the Multiclamp 700A amplifier closely matched the cancellation needed for most accurate recordings of amplitude and half-duration (minimal *χ*^2^) in the test circuit. With this hybrid voltage-/current-clamp approach (i.e. pipette capacitance neutralization in current-clamp mode using the value determined in voltage-clamp mode), optimal pipettes (5 pF), and an optimal amplifier (Multiclamp 700A) the errors in action potential amplitude and half-duration were smaller than 3% and 8%, respectively, making it feasible to reliably record presynaptic action potentials from small boutons.

### Action potentials in boutons of neocortical cultures and layer 5 neurons in brain slices have similar properties

Whole-bouton recordings in acute brain slices (Figure 2) were performed on an upright microscope. The capacitance of the recording apparatus in those recordings (glass pipette, pipette holder, and amplifier head-stage) was 6.46 ± 0.16 pF (n = 8; Figure 4A) which was significantly higher than the 5 pF found to be required for precise action potential recordings in the equivalent circuit (Fig 3). To reduce recording capacitance, we turned to an inverted microscope where immersion depth of the pipette is strongly reduced, which lowered recording capacitance to ~5.0 pF (n = 47; Figure 4A).

**Figure 4.**
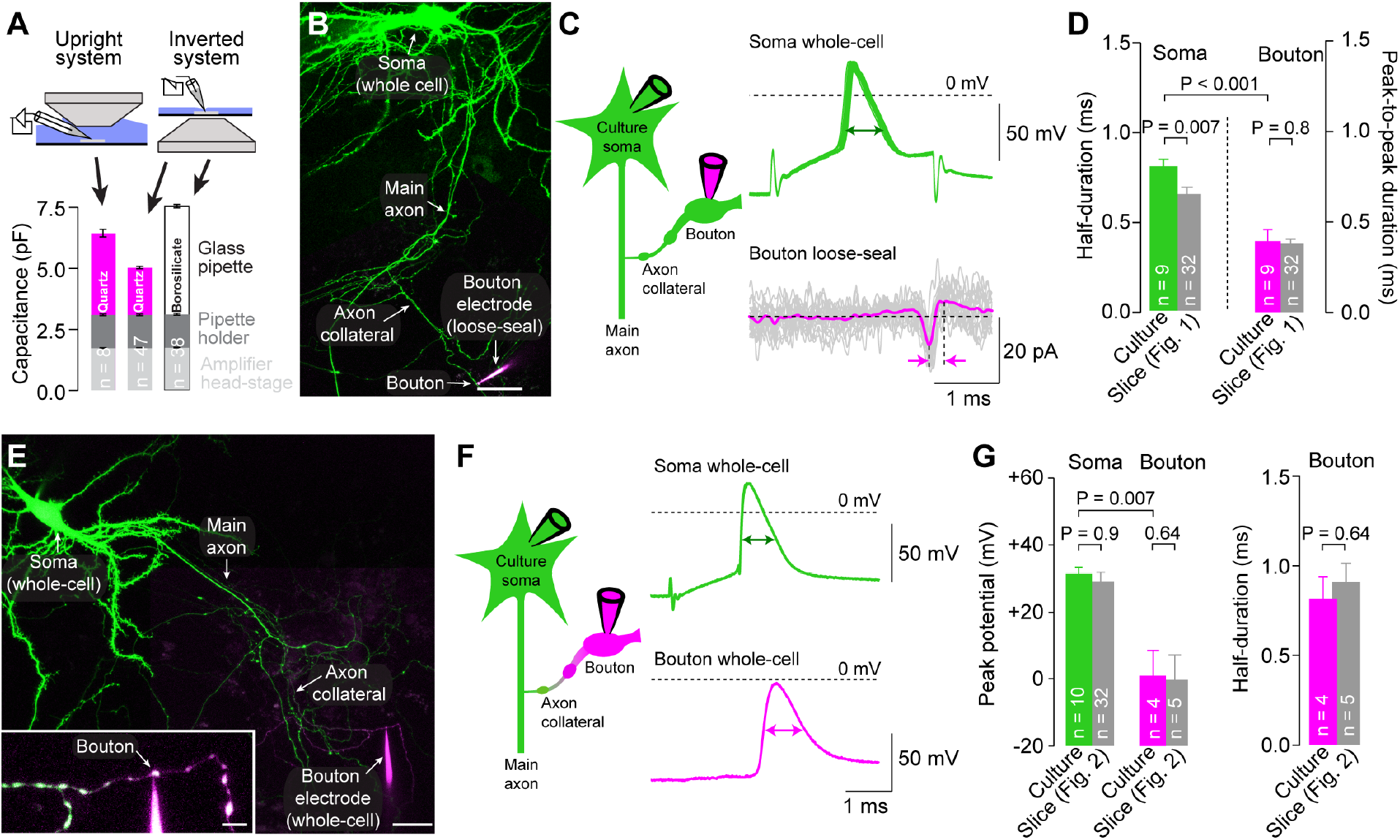
Action potential in boutons of neocortical cultures and layer 5 neurons in brain slices have similar properties. **(A)** Recording capacitance (glass pipette, pipette holder and amplifier head-stage capacitance) in recordings on an upright and inverted microscope with different glass pipettes (magenta: quartz glass pipettes; black: borosilicate glass pipettes; dark grey: pipette holder; light grey: amplifier head-stage). Bar graphs as mean ± SEM, n refers to the number of tested pipettes or iterations of capacitance compensations for amplifier head-stage and pipette holder. **(B)** 2P-image of a cultured excitatory neuron recorded in whole-cell mode and filled with 20 μM Atto-488 (green; maximum z-projection of a stack of 22 images, z-step size 1.0 μm). Action potential-evoked currents were simultaneously recorded from a bouton in loose-seal mode (magenta pipette, 100 μM Atto-594 contained in pipette). Scale bar 25 μm. **(C)** Left: Pictogram of the recording configuration. Right: Somatic action potentials (green) and action potential-evoked currents in a simultaneous loose-seal bouton recording (magenta). Action potential half-duration and current peak-to-peak duration indicated by arrows. **(D)** Half-duration of somatic action potentials (green) and peak-to-peak duration currents in boutons (magenta). Corresponding data from brain slice recordings are provided by grey bars for comparison (bars as mean ± SEM, n indicates number individual paired soma-bouton recordings). **(E)** 2P-image of a paired whole-cell bouton-soma recording (green pipette for somatic, magenta pipette for bouton recording) of a cultured excitatory neuron. Scale bar 25 μm, maximum z-projection of a stack of 21 images, z-step size 0.75 μm. *Inset:* Magnified whole-cell bouton recording, scale bar 5 μm, maximum z-projection of a stack of 18 images, z-step size 0.5 μm. **(F)** *Left:* Pictogram of the recording configuration. *Right:* Whole-cell recorded somatic (green) and bouton action potentials (magenta) upon somatic current-injection. Action potential half-durations indicated by arrows. **(G)** *Left:* Peak potential of somatic (green) and bouton (magenta) action potentials in paired bouton-soma recordings from cultured neurons. *Right:* Half-duration of somatic (green) and bouton (magenta) action potentials. Grey bars provide corresponding data from layer 5 pyramidal neuron recordings in brain slice for comparison (bar graphs as mean ± SEM, n indicates number of individual paired soma-bouton recordings).

The inverted microscope was incompatible with acute slice recordings necessitating the utilization of cultured neocortical neurons. Action potentials of boutons of 2-3 week-old cultured neurons were very similar to action potentials recorded from boutons of layer 5 pyramidal neurons in acute brain slices when recordings were made on an upright microscope. In these important control experiments, loose-seal action potentials of boutons were briefer than the simultaneously recorded somatic whole-cell spikes (Figures 4B – D) and the action potential half-width apparently increased in the whole-bouton configuration (Figures 4E – G). Notably, recorded parameters of action potentials in boutons were very similar in cultured neurons and layer 5 pyramidal neurons (grey bars in Figures 4D and G) consistent with the fact that the majority of boutons in the cultured neurons was excitatory (data not shown). This indicates that boutons of mature cultured neocritical neurons can serve as a valid model for the analysis of action potentials of excitatory boutons in the neocortex.

### Large and rapid action potentials in boutons of cultured neocortical neurons

To perform high-resolution current-clamp recordings from small boutons, we investigated action potentials in boutons of cultured neurons at an inverted microscope minimizing errors related to pipette capacitance. Bouton recordings were confirmed by (1) the presence of vesicle recycling detected by FM1-43 staining (Ryan et al., 1993; Sara et al., 2005; Smith et al., 2004), and (2) filling of the bouton and the adjacent axon with fluorescent internal solution contained in the recording electrode (Figure 5A).

**Figure 5.**
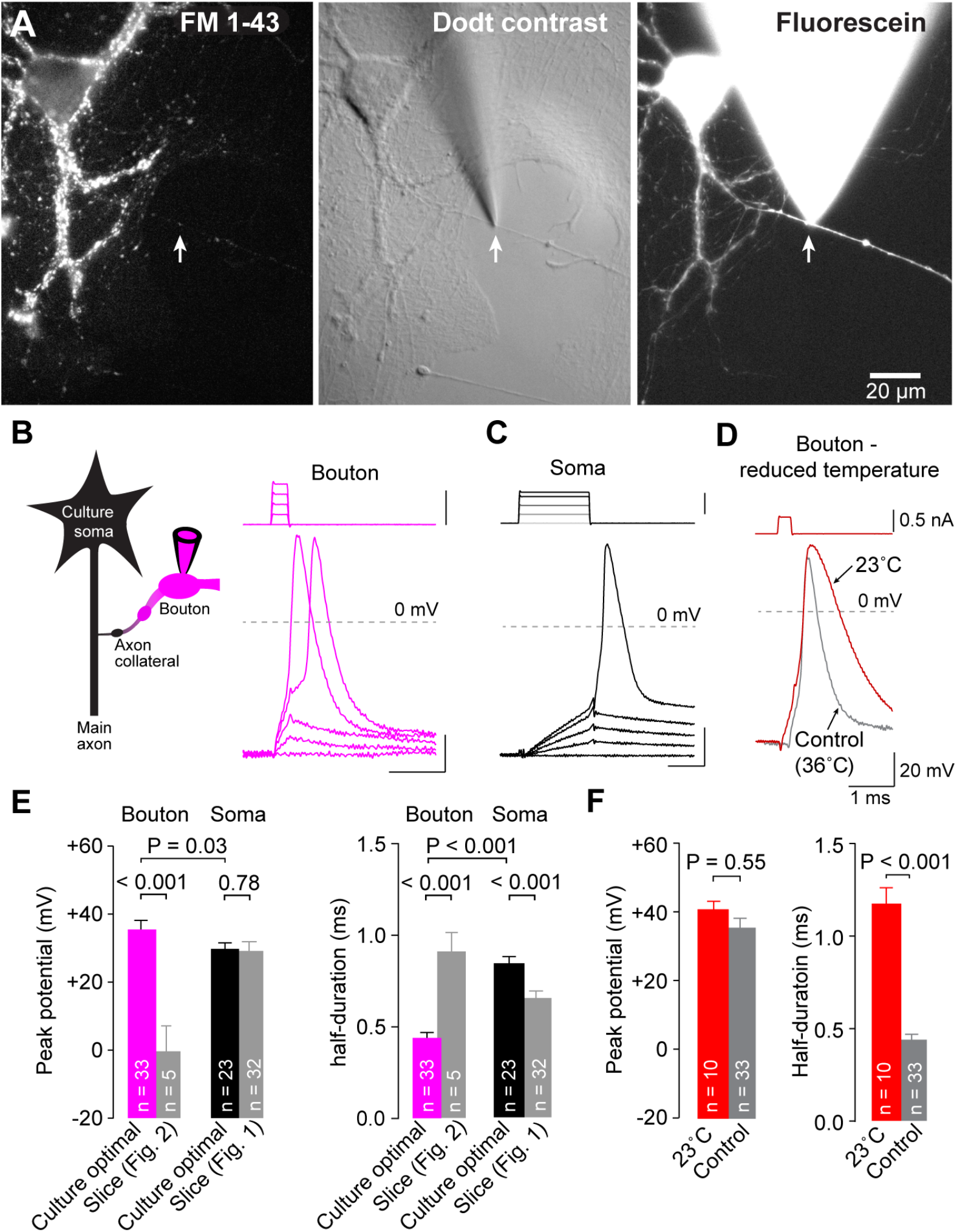
Large and rapid action potentials in boutons of cultured neocortical neurons. **(A)** Example image of a cultured neocortical cell: FM1-43 (*left*), and Dodt contrast (*middle*) and fluorescence image of a bouton whole-cell recording (*right*). **(B)** *Left:* Pictogram of the recording configuration. *Right:* Example action potentials evoked by current-injection in a whole-cell bouton recording with quartz glass pipettes on an inverted microscope (*right*). **(C)** Example somatic action potential evoked by current inj ection under the same conditions as in B. **(D)** Example bouton action potential recorded at reduced temperature (23 °C, red) compared to a control action potential recorded at 36 °C (grey, control action potential same as in B). **(E)** *Left:* Peak potential of evoked action potentials in boutons and somata of cultured neurons recorded with quartz glass pipettes on an inverted microscope. *Right:* Half-duration of evoked action potentials in boutons and somata. Grey bars indicate action potential peak potential and half-duration in bouton and soma recordings in brain slices on an upright microscope for comparison (bar graphs as mean ± SEM, n indicates number of individual paired soma-bouton recordings). **(F)** *Left*: Average peak potential of action potentials at reduced temperature compared to control condition. *Right*: Half-duration of action potentials at reduced temperature compared to control condition (color code as in D, bar graphs as mean ± SEM, n indicates number of individual bouton recordings). See also Figure S4.

In optimized whole-cell recordings from boutons in cultures, action potential half-duration was 452 ± 22 μs (n = 33; Figures 5B and E), far more consistent with the 380 μs value obtained with loose-seal recordings from boutons in slices. Furthermore, the bouton action potential peaked at 35.7 ± 2.7 mV (n = 33; Figures 5B and E), leading to action potential amplitudes of ~120 mV (117 ± 2.7 mV). Somatic action potentials were significantly broader and smaller contrasting with bouton action potentials (overshoot to 29.9 ± 1.7 mV and half-duration of 844 ± 35 μs; n = 23; P = 0.03 and P < 0.001, respectively; Figures 5C and E). Current injection used to locally elicit action potentials in boutons did not affect action potential properties, since spontaneous and evoked action potentials showed similar peak potentials and half-durations in recordings from boutons and somata under all experimental conditions (Figure S4).

As another test for the accuracy and the resolution of our optimized recordings in cultures, we recorded bouton action potentials at reduced temperature (23 °C; Figure 5D). Consistent with previous studies where temperature primarily impacted action potential duration and not amplitude (Borst and Sakmann, 1998; Schwarz and Eikhof, 1987), lowering temperatures broadened bouton action potentials by ~160 % (half-duration 1177 ± 83 μs, n = 10; P < 0.001) while amplitudes remained indistinguishable from those at physiological temperature (P = 0.55; Figure 5F).

Finally, we investigated how critical the pipette capacitance influences the action potential parameters. As predicted by our circuitry analysis (Figure 3), using borosilicate pipettes on an inverted microscope (~7.5 pF capacitance, Figure 4A) resulted in smaller and broader presynaptic spike (Figure S4E). Furthermore, pipette capacitance dithering distorted spikes measured at the bouton but not at the soma (Figure S4F). Taken together, optimized whole-cell recordings from boutons in cultures revealed large and brief presynaptic action potentials. Action potential duration was similar to loose-seal recordings from layer 5 pyramidal neurons, demonstrating that the discrepancy between loose-seal (Figure 1) and whole-cell recordings on the upright microscope (Fig 2) were due to distortion by larger recording capacitance.

### Presynaptic spike amplitude is independent of potassium channels and stable during synaptic scaling

K_v_1 channels have been proposed to contribute to plasticity of presynaptic action potentials by critically controlling action potential overshoot and voltage-gated calcium channel function in small boutons (Hoppa et al., 2014; Kawaguchi and Sakaba, 2015). The whole-bouton recording configuration in cultured neocortical neurons allowed us to directly test the hypothesis that action potential overshoot is controlled by potassium channel activity. Application of the selective K_v_1.1 and K_v_1.2 potassium channel blocker DTX-K (100 nM) strongly broadened presynaptic action potentials by ~60 % (half-duration 728 ± 55.6 μs, n = 12; P < 0.001) without affecting action potential overshoot (34 ± 4.6 mV, n = 12; P = 0.63; Figures 6A and B). Similar effects were observed in somatic recordings, where DTX broadened action potentials but did not affect action potential overshoot (data not shown). Even greater broadening of presynaptic action potentials was observed when K_v_1 and K_v_3 potassium channels were blocked by bath application of DTX-K and TEA (100 nM and 1 mM, respectively; half-duration 991 ± 113 μs, n = 8; P < 0.001; Figures 6A and B) while peak potentials were unaffected (35.5 ± 5.0 mV, n = 8; P = 0.67). These data indicate that in small cortical boutons, the action potential width but not peak voltage is regulated by K_v_1 and K_v_3 potassium channels.

**Figure 6.**
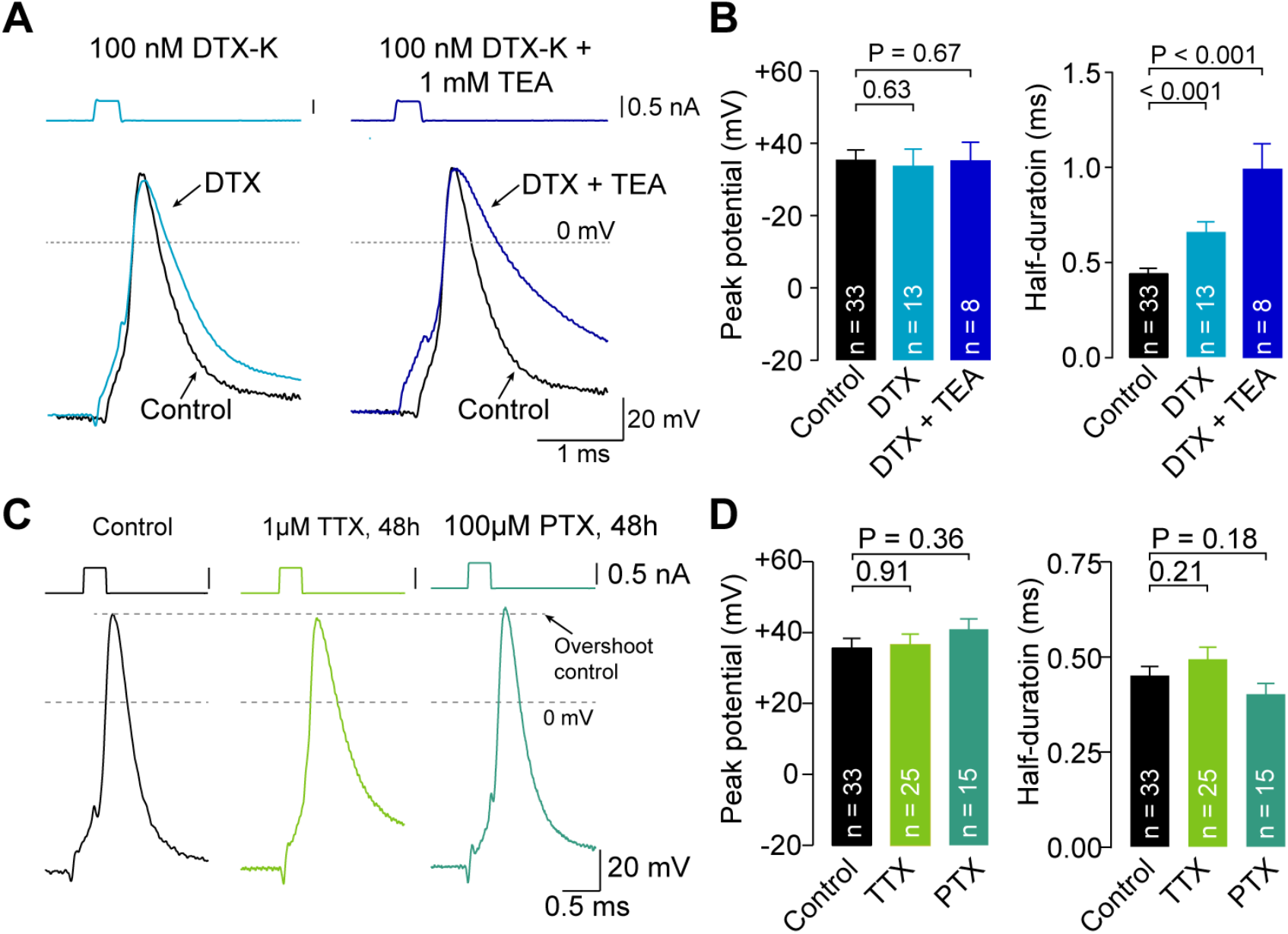
Presynaptic spike amplitude is independent of K_v_1 and K_v_3 potassium channels and stable during synaptic scaling. **(A)** *Left:* Example action potentials recorded from cultured boutons with 100 nM DTX-K included in the bath perfusion. *Right*: Example action potentials recorded from cultured boutons with 100 nM DTX-K + 1 mM TEA included in the bath perfusion. Representative control action potential recorded with quartz pipettes at physiological temperature (black, same as in Figure 5B) provided for comparison. Traces were aligned to the point of steepest rise during depolarization. **(B)** *Left:* Overshoot of action potentials recorded under the conditions illustrated in **a**. *Right:* Half-duration of action potentials recorded under the conditions illustrated in **a** (color code as in A, bar graphs as mean ± SEM, n indicates number of recordings from individual boutons). **(C)** Example action potentials recorded from cultured boutons under control conditions (same as in Figure 5B) and after 48h exposure to 1 μM TTX or 100 μM PTX (from *left* to *right*, respectively). **(D)** *Left:* Overshoot of action potentials recorded under the conditions illustrated in **c**. *Right:* Half-duration of action potentials recorded under the conditions illustrated in **c** (color code as in C, bar graphs as mean ± SEM, n indicates number of recordings from individual boutons, control data same as in Figure 5E). See also Figure S5.

Action potential waveform changes have also been hypothesized to mediate synaptic scaling or homeostatic synaptic plasticity (Li et al., 2020; Zhao et al., 2011). To test for changes in action potential waveform, we applied tetrodotoxin (TTX) or picrotoxin (PTX; 1μM and 100μM, respectively) for 48 hours prior to recordings, to increase or decrease synaptic strength, respectively (Murthy et al., 2001; Turrigiano et al., 1998; Zhao et al., 2011). As expected, TTX increased the excitatory quantal size after wash-off of the toxins (Figures S5B and C). However, neither intervention affected presynaptic spike amplitude (P = 0.91 and 0.36 for TTX and PTX, respectively) or half-duration (P = 0.21 and 0.18, respectively; Figures 6C and D). These results indicate that changes in bouton action potential waveform do not significantly contribute to homeostatic synaptic plasticity.

### Repeated stimulation evokes dynamic spike broadening in excitatory but not inhibitory boutons

To understand how the action potential waveform changes during short-term plasticity, trains of current injection-evoked action potentials were recorded in whole-bouton recordings of cultured neurons under optimal recordings conditions (Figure 7A; 20 action potentials at 20 or 50 Hz). The amplitude of the 1^st^ and 20^th^ action potential were similar at both frequencies (20 Hz: 98 ± 2%, n = 5, P = 0.44; 50 Hz: 97 ± 0.4%, n = 5, P = 0.13) whereas action potential half-duration increased profoundly over the same period (20 Hz: broadening to 138 ± 5%, n = 5, P = 0.06; 50 Hz: broadening to 136 ± 8%, n = 5, P = 0.06; Figure 7B).

**Figure 7.**
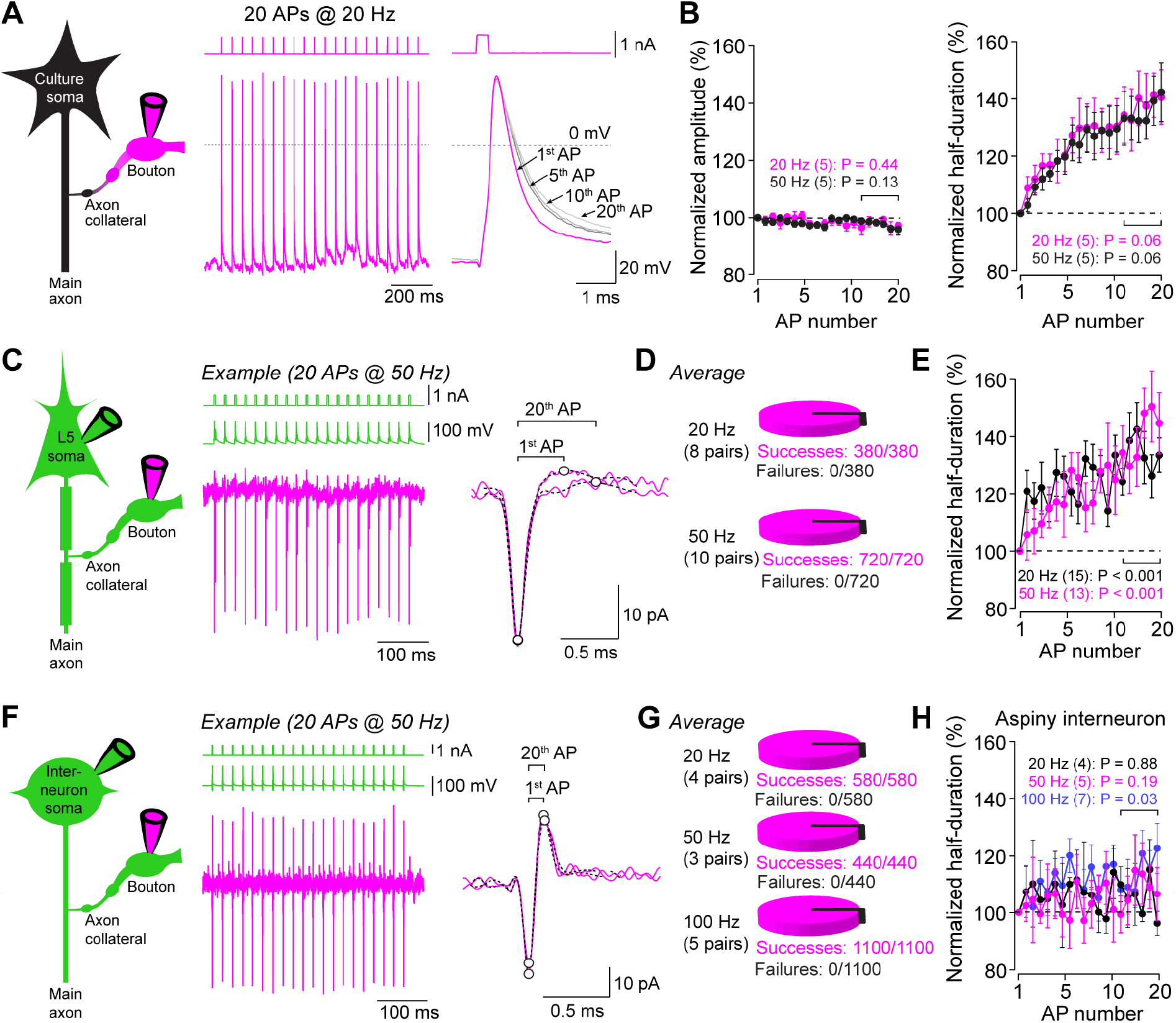
Repeated stimulation evokes dynamic spike broadening in excitatory but not inhibitory boutons. **(A)** *Left:* Pictogram of a whole-cell bouton recording in neocortical culture. *Middle:* Example action potential train (20 action potentials, 20 Hz) recorded from a bouton in culture. *Right:* Overlay of 1^st^, 5^th^, 10^th^, and 20^th^ action potential during train stimulation. **(B)** *Left:* Amplitudes of bouton action potentials in cultures normalized to the 1^st^ action potential during train stimulation at 20 Hz (black, n = 5) and 50 Hz (magenta, n = 5). *Right:* Half-duration *(right)* of bouton action potentials in neocortical cultures normalized to the 1^st^ action potential during the train. Brackets indicate last 5 train action potentials used for statistical analysis). **(C)** *Left:* Pictogram of a paired bouton-soma recording from a neocortical layer 5 pyramidal neuron. *Middle:* Example trace of a 50 Hz somatic train stimulation (green) and currents simultaneously recorded from the bouton in loose-seal mode (magenta). *Right*: Overlay of currents recorded in the bouton evoked by the 1^st^ and last train action potential. Circles indicate peaks of current components, brackets indicate peak-to-peak duration of currents. **(D)** Action potential propagation successes and failures during 20 Hz and 50 Hz train stimulation (380 and 720 action potentials in total, respectively). **(E)** Peak-to-peak duration of currents in bouton of layer 5 pyramidal neurons during 20 Hz (black, n = 15) and 50 Hz train stimulation (magenta, n = 13; durations normalized to 1^st^ current duration, bracket indicates last 5 train action potentials used for statistical analysis, n indicates the number of paired recordings). **(F)** *Left*: Pictogram of a paired bouton-soma recording from a neocortical interneuron. *Middle:* Example 50 Hz somatic train stimulation (green) and currents simultaneously recorded from a bouton in loose-seal mode (magenta). *Right*: Overlay of currents recorded in a bouton evoked by the 1^st^ and last train action potential. Circles indicate peaks of current components, brackets indicate peak-to-peak current durations. **(G)** Action potential propagation successes and failures during 20 Hz, 50 Hz, and 100 Hz train stimulation (580, 440, and 1100 action potentials in total, respectively). **(H)** Duration of action potential-evoked currents in bouton of neocortical interneurons during 20 Hz (black, n = 4), 50 Hz (magenta, n = 5), and 100 Hz train stimulation (blue, n = 7; durations normalized to 1^st^ current duration, bracket indicates last 5 train action potentials used for statistical analysis, n indicates the number of paired recordings).

In neocortical brain slices, we somatically evoked 20 Hz and 50 Hz action potential trains in layer 5 pyramidal neurons and recorded action potential-evoked currents from boutons in loose-seal mode (Figure 7C). Individual action potentials propagated reliably at both frequencies and no failures were observed at 20 Hz (n = 380 action potentials from 8 bouton-soma pairs) or 50 Hz (n = 720 action potentials from 10 bouton-soma pairs; Figure 7D). The peak-to-peak duration significantly increase during 20 Hz and 50 Hz action potential trains (20 Hz: 135 ± 6%, n = 15, P < 0.001; 50 Hz: 141 ± 9%, n = 13, P < 0.001; Figure 7E), similar to the spike broadening in cultured boutons. Action potential propagation into axon collaterals of neocortical fast-spiking interneurons was remarkably reliable and no failures were observed at frequencies up to 100 Hz (20 Hz: n = 580/4; 50 Hz: n = 440/3; 100 Hz: n = 1100/5 action potentials/bouton-soma pairs; Figures 7F and G). In contrast to recordings from layer 5 pyramidal neurons, only minor action potential broadening was observed in paired bouton-soma recordings from neocortical fast-spiking interneurons (Figure 7H). These data indicate stable amplitude, reliable conduction, and differential broadening of action potentials in small *en passant* boutons of layer 5 pyramidal neurons and interneurons during high-frequency transmission in the neocortex.

## DISCUSSION

Here, we report direct recordings of the spike amplitude and duration in conventional small *en passant* boutons of excitatory and inhibitory neocortical neurons. We extend previous studies (1) by establishing a hybrid voltage-/current-clamp approach to accurately measure action potentials from small cellular compartments; (2) by showing that the presynaptic action potential amplitude of small neocortical boutons is larger than at the soma, resistant to K_v_1 and K_v_3 potassium channel blockade, and unaffected by homeostatic long-term plasticity; and (3) by demonstrating differential spike broadening in excitatory and inhibitory neocortical boutons. Taken together these findings indicate that the neocortical presynaptic action potential is surprisingly robust and that presynaptic components of plasticity are mediated via changes in spike duration but not amplitude.

### Large presynaptic spikes at small neocortical boutons

Analysis of the action potential amplitude in small boutons of other brain regions revealed controversial results. In hippocampal cultured neurons, semiautomated targeted whole-cell patch-clamp recordings (Novak et al., 2013) revealed apparently small and DTX-*in*sensitive presynaptic spike amplitudes at room temperature (Vivekananda et al., 2017). Another study using genetically-encoded voltage sensor at the same preparation (Hoppa et al., 2014) also reported small but DTX-sensitive action potential amplitudes in boutons, providing a novel mechanism for presynaptic plasticity. However, large pipette capacitance might have limited amplitude resolution (cf. Figure 3) and the kinetics of the genetically-encoded voltage sensor (Bando et al., 2019; Maclaurin et al., 2013) predict underestimation of the action potential amplitude (see Figure S5A and legend), consistent with recent comparisons of genetically-encoded voltage sensors and electrophysiology (Gonzalez Sabater et al., 2020). In the cerebellum, direct recordings from small boutons of juvenile Purkinje neurons provide evidence for frequency-dependent attenuation of presynaptic spike amplitudes, which can be inhibited by potassium channel blockers (Kawaguchi and Sakaba, 2015). However, the resulting frequency-dependent synaptic depression was not observed in older animals (Turecek et al., 2016), consistent with DTX-*in*sensitive action potential amplitudes in small boutons of mature cerebellar basket cell interneurons (Begum et al., 2016). Finally, our results are consistent with large action potential amplitudes measured in thin axons of interneurons in the hippocampus (Hu and Jonas, 2014; Hu et al., 2018), and thin dopaminergic axons in the striatum (Kramer et al., 2020). Our data thus reveal a large presynaptic spike amplitude in the neocortex and resolve the controversy regarding the spike amplitude of small excitatory boutons.

The amplitude of presynaptic action potentials in small boutons reported here agrees well with that of large boutons (Alle et al., 2011; Geiger and Jonas, 2000; Ishikawa et al., 2003; Ritzau-Jost et al., 2014; Taschenberger and von Gersdorff, 2000). However, evidence for potassium channel sensitivity of the spike amplitude has been described at the calyx of Held synapse (Ishikawa et al., 2003), where potassium channel facilitation was shown to attenuate spike amplitudes during high-frequency firing (Yang et al., 2014). Furthermore, the action potential amplitude increases upon blockade of K_v_1-channels (Kole et al., 2007; Shu et al., 2007) in the axon initial segment of layer 5 pyramidal neurons where the K_v_1 channel density was reported to be highest (Kole et al., 2007). These data suggest that amplitude of the action potentials in some large boutons and the axon initial segment is limited by potassium channel activation but at the majority of boutons, particulariy the here studied, prototypical small *en passant* boutons in the neocortex, action potential amplitude is large and not limited by potassium channel activation.

### Duration of presynaptic spikes

We found rapid action potentials with a half-duration of 0.45 ms and 0.23 ms at 36°C in small excitatory and inhibitory neocortical boutons, respectively. These values appear consistent with some studies on small excitatory and inhibitory boutons in other brain regions using presynaptic patch clamp recordings (Hu and Jonas, 2014; Vivekananda et al., 2017) and recordings with fast voltage-sensitive dyes (Casale et al., 2015; Foust et al., 2011) although comparison is complicated by differences in recording temperature and the definition of the half-duration. In our recordings from cultured neocortical neurons, selectively blocking K_v_1.1 and 1.2 channels (DTX-K) or blocking K_v_1 and K_v_3 channels (DTX-K and TEA) substantially broadened presynaptic action potentials by ~54 % and ~119 %, respectively (Figure 6). This is consistent with a series of studies demonstrating the contribution of K_v_1 channels to the repolarization at boutons of layer 5 pyramidal neurons and fast-spiking interneurons in the neocortex (Casale et al., 2015; Foust et al., 2011), small boutons of cultured hippocampal neurons (Vivekananda et al., 2017), small boutons of cerebellar basket (Begum et al., 2016; Southan and Robertson, 1998), small boutons of cerebellar stellate cell (Rowan et al., 2016; Rowan et al., 2014), and large central boutons (Alle et al., 2011; Ishikawa et al., 2003; Ritzau-Jost et al., 2014). There is also accumluating evidence for K_v_3 channel-dependent control of action potential duration in small (Hoppa et al., 2014; Rowan et al., 2014) and large boutons (Alle et al., 2011; Ritzau-Jost et al., 2014; Wang and Kaczmarek, 1998). This is notably different from the potassium channel independent repolarization in peripheral nodes of Ranvier (Kanda et al., 2019). Thus, our data indicate that K_v_1 and K_v_3 channels cooperatively regulate the action potential duration in small cortical boutons.

### Current-clamp recordings from small cellular structures

The standard approach for current-clamp recordings is the gradual increase of the pipette capacitance compensation until ringing of the potential occurs (Marty and Neher, 2009). At small cellular structures, this classical approach was unreliable resulting in errors in the amplitudes of action potentials (Figure S3). Other approaches for pipette capacitance compensation in the current clamp mode have been described (Riedemann et al., 2016), but have not been tested for small cellular compartments investigated here (<1 pF). We therefore established a hybrid voltage-/current-clamp approach using a test circuit to mimic a small bouton and systematically compared the accuracy of patch-clamp amplifiers (Figure 3). Although the problems arising from the combination of voltage- and current-clamp circuits in one amplifier (Magistretti et al., 1996) have been solved by most modern amplifiers, permitting adequate voltage- and current-clamp recordings under a wide range of conditions, a systematic comparison of the current-clamp performance of commonly used amplifiers is lacking. We observed substantial differences between different amplifiers (Figure 3) when recording directly at small boutons, where the capacitance is relatively low compared to that of the recording apparatus. Furthermore, we show that, even with optimal pipette capacitance cancellation, the absolute size of pipette capacitance limits the temporal resolution of current-clamp recordings. Finally, our results show that not only the measured voltage (V_cc_), but in addition the membrane potential itself (V_m_) can be dramatically altered by the pipette capacitance (Figure 3B). Rapid presynaptic action potentials are particularity affected by these errors but the high-frequency component of postsynaptic potentials measured with dendritic recordings are expected to be equally distorted (Davie et al., 2006; Jayant et al., 2017). We suspect that these errors are often underestimated, although they can be comparable to well-established large clamping errors during voltage clamp (Williams and Mitchell, 2008). Yet, the here-described hybrid voltage-/current-clamp approach, in combination with optimal amplifiers and low-capacitance pipettes, allows reliable measurements of action potentials with a temporal resolution of <25 μs and an amplitude resolution of <3% at small subcellular structures.

### Differential broadening and plasticity of the presynaptic spike

Spike duration increased at excitatory but not inhibitory neocortical boutons during high-frequency firing (Figure 7). This finding is consistent with other studies at excitatory and inhibitory bouton synapses in the hippocampus (Geiger and Jonas, 2000; Hu and Jonas, 2014; Ma et al., 2017) and emphasizes distinctions in the regulatory pathways of excitatory and inhibitory synapses in addition to neurotransmitter identity (Tsintsadze et al., 2017). The minimal spike broadening in boutons of fast-spiking interneurons is consistent with K_v_3 channels localized at boutons of fast-spiking interneurons in the neocortex (Goldberg et al., 2005), which have been shown to suppress spike broadening at the soma of hippocampal interneurons before (Lien and Jonas, 2003). The differential presynaptic spike broadening will contribute to the short-term plasticity of excitatory and inhibitory transmission and therefore dynamically affect the excitation/inhibition (E/I)-ratio during high-frequency excitatory and inhibitory inputs (Galarreta and Hestrin, 1998; Xue et al., 2014).

The reliability of the neocortical presynaptic spike shape during other forms of neuroplasticity was surprising. In particular, the insensitivity of the presynaptic spike duration to periods of reduced activity (Figure 6) was remarkable since somatic action potential duration is substantially increased by maneuvers that promote synaptic scaling (Li et al., 2020), indicating that the pathways by which BK-channel-dependent spike broadening occurs following synaptic scaling are effectively confined to neuronal compartments upstream of the axon. The presynaptic mechanisms of synaptic scaling in the neocortex thus differ from depolarization-induced potentiation of excitation at large hippocampal mossy fiber boutons mediated by spike broadening (Carta et al., 2014). In summary, in neocortical boutons the spike amplitude is remarkably constant and dynamic changes are confined to the spike duration of excitatory neurons.

## METHODS

### Ethics

All animal procedures were in accordance with the European (EU Directive 2010/63/EU, Annex IV for animal experiments), national and Leipzig University guidelines as well as the U.S. Public Health Service Policy on Humane Care and Use of Laboratory Animals and the National Institutes of Health Guide for the Care and Use of Laboratory Animals. All animal procedures were approved in advance by the institutional Leipzig University Ethics Committees, the federal Saxonian Animal Welfare Committee as well as the V.A. Portland Health Care System Institutional Animal Care and Use Committee.

### Animal models

All experiments were performed using postnatal day 0 or 1 (for neocortical cultures) or postnatal day 17 – 32 C57BL/6N mice (for brain slice recordings) of either sex. Adult mice and mother animals were fed *ad libitum*.

### Preparation of acute neocortical brain slice

Mice were anesthetized by isoflurane (2.5% in 4l oxygen/min) and rapidly killed by decapitation. Brain hemispheres were mounted in a chamber filled with chilled ACSF, sliced coronally into 300 *μ*m thin slices (VT1200 vibratome, Leica Microsystems, Wetzlar, Germany), and incubated in ACSF at 37 °C for 30 minutes after slicing. Afterwards, slices were transferred to ACSF at room temperature until usage.

### Preparation of neocortical cell cultures

P0 or P1 mice were decapitated and the cerebral cortices were removed. For subsets of the data shown in Figures 5 and 7 that were conducted in Portland, Oregon, USA, decapitation was performed following general anesthesia with isoflurane. The neocortices were dissected in icecold Hank’s balanced saline solution and cut into 5 – 8 pieces per hemisphere. Brain pieces were digested by Trypsin (5 mg, dissolved in digestion solution; for composition see below) for 5 minutes at 37 °C. Afterwards, trypsin was stopped by ice-cold standard medium and the supernatant was discarded before pieces were enzymatically and mechanically dissociated in standard medium containing DNAse (10 mg/ml) by trituration using Pasteur pipettes of narrowing tip diameters. Subsequently the supernatant was separated and centrifuged. After resuspension in standard medium, vital (Trypan blue negative) cells were counted and the suspension was plated onto coverslips (50×10^3^ vital cells/coverslip) coated with Matrigel in 24 well plates. Forty-eight hours after plating, the medium was fully replaced by standard medium containing 4 μM cytosine arabinoside in order to limit glial growth. Ninety-six hours after plating, the medium was again fully replaced by standard medium. Cells were incubated at 37 °C, 93% humidity, and room air plus 5% CO_2_ until they were used following 14 to 19 days in culture. Standard medium was prepared from 1l MEM (Earle’s salts + L-Glutamine) mixed with 5g Glucose, 0.2g NaHCO_3_, 0.1g bovine holo-Transferrin, 0.025g bovine insulin, 50 ml fetal bovine serum and 10 ml B-27. Digestion solution contained the following (in mM): 137 NaCl, 5 KCl, 7 Na_2_HPO_4_, 25 HEPES, pH adjusted to 7.2 by NaOH.

### Electrophysiology

Voltage- and current-clamp recordings from boutons as well a subset of somatic recordings (Figure 5 and Figures S1 and S4) were performed with quartz glass pipettes using a Multiclamp 700A patch-clamp amplifier (Molecular Devices, San Jose, USA). Quartz glass pipettes (without filament; Heraeus Quartzglas, Kleinostheim, Germany) were pulled with a DMZ Universal Electrode Puller with oxygen-hydrogen burner (Zeitz Instruments, Martinsried, Germany; Dudel et al., 2000). In the majority of somatic recordings, somata were recorded using borosilicate glass pipettes and a HEKA EPC10/2 amplifier (HEKA Elektronik, Lambrecht/Pfalz, Germany). Borosilicate glass pipettes were pulled from borosilicate glass (Science Products, Hofheim, Germany) by a DMZ Universal Electrode Puller with filament heating (Zeitz Instruments, Martinsried, Germany).

Pipette resistances were 3 – 7 MΩ and 7 – 15 MΩ for somatic and small bouton recordings, respectively, when filled with a potassium gluconate-based internal solution. Membrane potentials were corrected for the calculated 12 mV liquid junction potential. Unless otherwise noted, recordings were performed at physiological temperature (36 ± 1 °C). Pipette capacitance was systematically minimized by the use of shortest possible pipettes (~1.5 and ~4 cm length for recordings on inverted and upright microscopes, respectively), low bath perfusion levels, and the application of silicone grease onto the recording electrode wire to prevent internal solution from being pulled up by adhesive forces. Pipette coating did not further reduce pipette capacitance in our hands, potentially due to the low bath perfusion levels achieved on the inverted microscope. Pipettes were fixed on a custom-built pipette holder mounted on a micromanipulator (Kleindiek Nanotechnik, Reutlingen, Germany). All current-clamp recordings were filtered with the internal 10 kHz 4-pole Bessel filter of the Multiclamp 700A amplifier or the internal 10 kHz 8-pole Bessel filter of the HEKA EPC10/2 amplifier and subsequently digitized (200 kHz) with the HEKA EPC10/2 using Patchmaster software (HEKA Elektronik, Lambrecht/Pfalz, Germany).

For pipette capacitance compensation during whole-cell bouton recordings, we used a hybrid voltage-/current-clamp approach validated with an electrical circuitry (Figure 3, see main text and below). Whole-cell configuration was achieved by applying short repetitive pulses of positive and negative pressure applied via a hand-operated syringe coupled to the custom-build pipette holder. After establishing whole-cell configuration and automatic determination of the pipette capacitance (including amplifier head-stage and pipette holder), the amplifier was changed to current-clamp mode and the pipette capacitance was cancelled using the capacitance value determined in voltage clamp mode. For recordings with the Multiclamp 700A amplifier, the slow pipette capacitance compensation was not used since it did not affect action potential shape. The 500 MΩ feedback resistor of the head-stage circuitry of the Multiclamp 700A amplifier was used during current-clamp recordings to optimally match expected load resistance during our experiments within the 1:5 to 5 fold range recommended by the manufacturer. Holding currents were adjusted to hold membrane potentials at −80 mV (liquid-junction potential corrected). To investigate the potential effect of the resting membrane potential on the action potential amplitudes, we varied the holding potential in a subset of experiments (n = 17 bouton recordings). An exponential fit to the population data of peak potentials over holding potentials predicted an action potential peak at 30.3 mV for −60 mV holding potential with less than 7.4 mV change in peak potential for holding potentials between −90 and −60 mV. Bridge compensation was applied subsequent to capacitance neutralization and adjusted to minimize the current injection artifact.

### Recordings from acute neocortical brain slices

Brain slices were recorded on an upright two-photon laser scanning microscope (Femtonics, Budapest, Hungary) equipped with a pulsed Ti:Sapphire laser (MaiTai, SpectraPhysics, Santa Clara, USA) tuned to 805 nm and a 60x/1.0 NA objective (Olympus, Shinjuku, Japan). Internal recording solutions contained 100 – 200 μM Atto-594 Carboxy (ATTO-TEC, Siegen, Germany) for bouton recordings and 20 μM Atto-488 Carboxy (ATTO-TEC, Siegen, Germany) for somatic recordings to confirm successful whole-cell recordings and visualize the axonal arborization, respectively. Boutons were recognized as local swellings of ~1 μm diameter on axonal collaterals. For loose-seal recordings, quartz pipettes were placed on the bouton and the resulting pipette resistance did not exceed 100 MΩ.

### Recordings from cultured neocortical neurons

Cell cultures were mounted on an inverted (Nikon TI-U, Nikon Instruments, Melville, USA or Scientifica SliceScope, Scientifica, Uckfield, United Kingdom) visualized by a 100x or 60x water-immersion objective (NA 1.1 or 1.0, respectively) and difference interference or Dodt contrast optics. For cultures recorded on the upright microscope (Figure 4), recordings were performed similar to brain slices recordings (see above). As an alternative to prior somatic filling with fluorescent internal solution, boutons were recognized as fluorescent puncta stained by depolarization-dependent endocytosis using FM1-43 (10 μM in isotonic Tyrode variant containing 90 mM KCl for 90 s). These hotspots coincided with axonal varicosities. With more experience, axonal varicosities were reliably detected as FM1-43-positive boutons. In a subset of recordings boutons were therefor visually identified without prior FM1-43 staining. In all cases, axonal origin was confirmed by the filling of the bouton and the adjacent axon with fluorophore (20 μM Atto-488 Carboxy; ATTO-TEC, Siegen, Germany) contained in the pipette solution once whole-cell configuration was attained and there were no systematic differences between boutons with and without FM1-43 pre-staining.

### Passive parameters of boutons

Presynaptic bouton passive parameters were measured in cell-attached mode and immediately after establishing whole-cell mode using a −10 mV step (5 ms). At least one hundred trials were averaged for each recording condition and cell-attached currents were subtracted from wholecell currents (Figure S3A). Remaining currents were analyzed with a two-compartment model as previously described (Hallermann et al., 2003) providing the capacitances of both compartments (C_1_ and C_2_).

### Analysis of action potentials

Data were analyzed using IGOR PRO software (WaveMetrics, Lake Oswego, OR, USA; Version 6.32A), the Patcher’s Power Tool (Version 2.19 Carbon, 15.03.2011) and NeuroMatic (Version 2.00, 15.09.2008; Rothman and Silver, 2018) extensions, as well as self-written IGOR PRO scripts. To exclude the effect of mis-balanced bridge compensation in bouton recordings, only action potentials that occurred after the end of the second current injection artifact (elicited by the end of current-injection) were analyzed. For somatic recordings, current injection artifacts were blanked over 50 μs at the beginning and the end of the current injection and only experiments in which blanking did not affect action potential amplitude and half-duration were subsequently analyzed.

Action potential amplitudes were measured from the average membrane resting potential between 15 and 5 ms before action potential initiation to the action potential peak voltage.

Action potential half-durations were calculated as the duration at half-maximal action potential amplitude. In a subset of recordings, action potential initiation occurred after the end of the current injection. In those experiments, the action potential threshold was determined as first derivative of voltages crossing 50 V s^-1^ (Kole and Stuart, 2008) and action potential amplitudes were calculated from threshold and the half-durations was calculated as the duration at the half of the corresponding amplitude. The temperature coefficient (Q_10_) of action potential half-durations was 2.2 and was calculated as Q_10_ = (D_2_/D_1_)^10°C/ΔT^ from the action potential half-duration at physiological temperature (D_2_) and room temperature (D_1_) and ΔT = 13 °C = 36 – 23 °C.

### Analysis of currents from loose seal recordings

Loose-seal currents were analyzed to derive action potential durations by self-written Python routines. In short, currents were digitally filtered (7.5 to 15 kHz, 8-pole Bessel filter), interpolated and further filtered with a smooth-spline interpolation algorithm, alignment to the negative peak of the interpolated current (representing the sodium current), and averaged. The peak-to-peak spike duration was the interval between the negative and positive peaks representing peak sodium and potassium currents. For action potential conduction failure analysis, the negative sodium current peak was determined in a 0.5 ms time window at the interpolated individual traces. As a control, noise peaks were determined in identical ways in each trace in a window from 1.5 to 1 ms before the beginning of the sodium current windows. Noise and sodium peak histograms were fitted with double gaussian fits according to

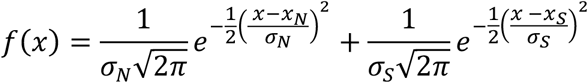

where *x_N_* and *x_s_* denote the mean of the noise and sodium peaks, respectively, and *σ_N_* and *σ_s_* denote the standard deviation of the noise and sodium peaks, respectively.

Only experiments with sufficient signal-to-noise ratio were analyzed in which the overlap of both gaussians was < 10^-4^. To assess the reliability of the detection of action potentials, the following two probabilities were determined for each experiment: The probability *P*_1_ of finding a noise peak larger than the smallest sodium peak, *S_smallest_*, in the distribution of the noise peaks according to

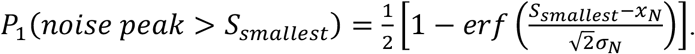

The probability *P*_2_ of finding a sodium peak smaller than the largest noise peak, *N_largest_*, in the distribution of the sodium peaks according to

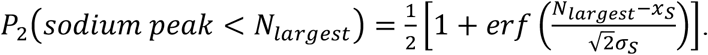

On average, *P*_1_ and *P*_2_ were 2.6×10^-7^ and 4.4×10^-6^, respectively, in 16 paired loose-seal bouton and whole-cell soma recordings. *P*_1_ and *P*_2_ were always < 10^-4^, indicating that the chance of false-positively detecting an action potential, when in fact a conduction failure had occurred, was always well below 1 in 10.000. Thus, our detection reliability was sufficient do distinguish between successful action potential conduction and conduction failure.

### Comparison of borosilicate and quartz glass pipettes

Light microscopic images of pipettes were taken with an Olympus CX41 light microscope (Olympus, Tokyo, Japan) at 10x and 100x (LMPlanFL N objective, NA 0.8; Olympus, Tokyo, Japan) magnification. For electron microscopy, the individual pipettes were placed in brass pipette holders and fixed with conducting silver cement (EMS, Hatfield, USA). Next, pipettes were sputter-coated with platinum in an upright position and under permanent rotation (Quorum, Laughton, UK). The final thickness was monitored and adjusted to 5 nm with a film thickness monitor. Electron microscopy was performed using a Zeiss SIGMA electron microscope (Zeiss, NTS, Oberkochen, Germany). Images were acquired at 5 kV using an Inlens detector and an aperture of 10 μm to minimize negative charges on the specimen. For image acquisition, the resolution was adjusted to pixel sizes of 0.3 – 0.7 nm.

The pipette tip opening area (a) was quantified manually in FIJI (Schindelin et al., 2012) and the pipette tip radius (r) was calculated by 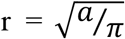. For the quantification of pipette capacitances (C_pip_) and pipette resistances (R_pip_), sister pipettes of those imaged by electron microscopy were placed under realistic recording conditions (holder and bath perfusion as during electrophysiological recordings) and the capacitance of the recording apparatus and pipette resistance were determined by built-in routines of the Multiclamp 700A amplifier in voltage-clamp mode.

The capacitance of glass pipettes, pipette holder and amplifier head-stage contributed to overall parasitic capacitance. Glass pipette capacitance was calculated by subtracting the capacitance of the amplifier’s head-stage and of the pipette holder from the overall capacitance. Capacitances of quartz glass pipettes alone were 3.35 ± 0.16 pF (n = 8) and 1.95 ± 0.05 pF (n = 47) on the upright and inverted microscope, respectively. Capacitance of borosilicate glass pipettes alone was 4.44 ± 0.07 pF on an inverted microscope. Capacitance of the Axon Multiclamp 700A amplifier head-stage was 1.74 ± 0.01 pF and determined by automated capacitance cancellation before connecting the pipette holder. Pipette holder capacitance was 1.37 ± 0.04 pF and calculated by subtracting the head-stage capacitance from the capacitance measured with holder and head-stage connected. Both, amplifier head-stage and pipette holder capacitance were recorded 10 times at the gain used in the subsequent experiments.

### Electronic equivalent circuit model

An electronic equivalent circuit model (Figure 3A) was developed to investigate the impact of pipette capacitance (C_pip_) and C_pip_ cancellation on the ability of different amplifiers to measure action potential parameters correctly. Voltage commands (V_in_) were applied via a resistor branch of 10 series-connected 100 MΩ resistors (1 GΩ total resistance, resistors in series to reduce total stray capacitance) to an on-board operational amplifier (OpAmp). Although we have used an operational amplifier with a low input capacitance of ~1 pF (LTC6268IS8#PBF, Analog Devices, Norwood, Massachusetts, USA), the capacitance of the equivalent circuit representing the bouton was ~2 pF. Because this is larger than the bouton and the adjacent axonal compartment, we electrically neutralized part of the input capacitance to reach ~0.5 pF (representing the smallest sum of C_1_ and C_2_ in our recordings; Figure S3A) by a circuit providing current to the input from the amplifier output via a feedback capacitor (this is equivalent to the capacitance neutralization circuits found in amplifiers). Voltages were recorded at two independent voltage outputs: One connected directly to the OpAmp output, which was treated as recording the real voltages of the small bouton compartment (V_m_). A second output was separated from the OpAmp by a 10 MΩ resistor chain (10 connected 1MΩ resistors) with either ~5 pF or ~7.5 pF stray capacitance, thus electronically resembling a recording pipette (‘pipette branch’), and was recorded by different current-clamp amplifiers (V_cc_). We did not systematically study the effect of different series resistance in our circuitry analysis because we realized that the series resistance had little impact on the action potential amplitude. For example, an increase in the series resistance from ~15 to ~50 MΩ changed the peak action potential amplitude by <5% (n = 4 bouton recordings).

### Amplifier comparisons

The following amplifiers were tested systematically by the electronic equivalent circuit model: Axon Instruments Multiclamp 700A (Molecular Devices, San Jose, USA), HEKA EPC10/2 (HEKA Elektronik, Lambrecht/Pfalz, Germany), NPI BA-03X and NPI ELC-03XS (both npi electronic, Tamm, Germany). The impact of pipette capacitance and capacitance cancellation on action potentials was investigated as follows (Figure 3B): First, the pipette branch was separated from the circuit (R19 in Figure 3A disconnected) and voltage commands were adjusted to reveal action potential-like voltage transients at V_m_ (termed ‘target’). After reconnecting the pipette branch, voltages at V_m_ and V_cc_ were recorded in current-clamp mode while systematically changing the capacitance compensation value of the respective amplifiers. Increasing capacitance compensation generally increased amplitudes and shortened durations of recorded action potentials, approaching and in some cases exceeding the ‘target’ (Fig 3B). For each capacitance cancellation setting, amplitude and half-duration of recorded action potentials were quantified and compared to the ‘target’ waveform (horizontal dashed lines in Figures 3C and D). Deviations of the recorded action potentials from the ‘target’ were furthermore determined by the sum of squared errors (*χ*^2^; calculated over 5 ms following the start of voltage commands; Figure 3C). Additionally, exact capacitance of the isolated lower-capacitance pipette branch (~5 pF) was determined by each combined voltage-clamp/current-clamp amplifier individually (all amplifiers but NPI BA03X, which is a pure current-clamp amplifier) in voltage-clamp mode using the built-in C_pip_ cancellation routines (Multiclamp 700A: 5.2 pF; HEKA EPC10/2: 4.6 pF; NPI BA-03X: 5 pF; see also vertical dashed lines in Figure 3D). Capacitance compensation settings leading to the optimal match between recorded and the ‘target’ action potential were compared to capacitances determined for pipette branches by the same amplifier.

Of note, the NPI BA-03X amplifier is a pure current-clamp amplifier and does not provide voltage-clamp mode, precluding determination of pipette branch capacitance under our experimental conditions. For the NPI ELC-03XS amplifier, the pipette capacitance automatically determined in voltage-clamp mode could not be transferred to current-clamp mode due to voltage oscillations occurring already at substantially lower capacitance cancellation. The HEKA EPC10/2 amplifier automatically reduces pipette capacitance cancellation by 0.5 pF upon switching from voltage-clamp to current-clamp mode. We accounted for this by manually adding 0.5 pF to the automatically determined capacitance cancellation in voltage-clamp mode before switching to current-clamp mode. We also repeated these analyses with shorter action potential-like commands (half-duration ~170 μs) and 10 ms square voltage commands revealing similar results but stronger deviations from the ‘target’ in the case of shorter action potential-like commands (data not shown).

### Limitations of the hybrid voltage-/current-clamp approach

In principle, compensation for stray capacitances determined in voltage-clamp mode during current-clamp recordings could introduces the two following systematic errors. (1) During transition from cell-attached to whole-cell the capacitance of the membrane forming the gigaseal is lost. However, the capacitance of this patch of membrane was calculated for an average 10 MΩ quartz glass pipette (opening radius 220 nm; cf. Figure S2C) and the estimate of 3 fF and 12 fF accounted for 1 % and 4 % of the total membrane area of the small compartment (C_1_ = 0.29 pF; Figure S3A) when assuming a membranous hemisphere directly at the pipette tip or a two times larger hemisphere inside the pipette lumen, respectively. This error is thus negligible. (2) The capacitance of the head-stage will change upon switching from voltage-to current-clamp. However, our analysis of the equivalent circuit (Figure 3) demonstrated that this error is for example 0.1 pF for the Multiclamp 700A amplifier and without explicit correction for this error the amplitude and the half-duration could be determined with an error smaller than 3-8%, respectively.

### Mathematical model of the voltage sensitive dye

The fluorescence response of the genetically-encoded voltage sensor was predicted with simple exponential rise and decay during mock action potentials (rectangular action potentials with average amplitudes and half-durations predicted from our recordings). The exponential rise and decay time constants were 0.4 and 0.6 ms, respectively, as reported for Archaerhodopsin (Bando et al., 2019; Maclaurin et al., 2013; time constants for steps from −70 to +30 mV and from +30 to −70 mV, respectively). Durations of mock action potentials at 30 °C were derived from recorded action potential durations at 36 °C using the here-determined Q_10_ of 2.2 for the temperature dependence of the action potential half-duration.

### Solutions and reagents

For recordings from cell cultures and brain slices on the upright microscope (Figures 1, 2, 4, and 7), extracellular artificial cerebro-spinal fluid (ACSF) contained (in mM): 125 NaCl, 3 KCl, 25 Glucose, NaH_2_CO_3_ 25, Na_2_HPO_4_ 1.25, 1.1 CaCl_2_, 1.1 MgCl_2_. For recordings from cultured neurons on the inverted microscope (Figures 5 – 7), extracellular Tyrode-based solution contained (in mM): 150 NaCl, 4 KCl, 10 HEPES, 10 Glucose, 1.1 CaCl_2_, 1.1 MgCl_2_, 0.01 SR95531, pH adjusted to 7.35 by NaOH. The sodium reversal potential for the used solutions was +71 mV. For pharmacological block of voltage-activated potassium channels (Figure 6), 100 nM DTX-K (Alomone Labs, Jerusalem, Israel) or 100 nM DTX-K and 1 mM TEA-Cl (final concentrations) were added to the external solution. For induction of homeostatic plasticity, either 1μM TTX or 100 μM PTX were applied to the incubating cell culture medium 48 – 56 hours before recordings were performed. Recording pipettes were filled with internal solution containing (in mM): 150 K-gluconate, 3 Mg-ATP, 0.3 Na-GTP, 10 K-Hepes, 10 NaCl and 0.2 EGTA, pH adjusted to 7.2 by KOH. Chemicals and toxins were obtained from Sigma-Aldrich (St. Louis, MO, USA) unless otherwise noted.

### Statistics

Data are expressed as mean ± SEM. The indicated sample sizes (n) are provided as the number of individual somatic/presynaptic recordings. All data sets obtained from cultured neurons were from ≥ 3 independent cell cultures. A systematic variation between cell independent cultures was not observed. All experimental groups were independent and tested for statistical difference by the Mann-Whitney *U* test in IGOR PRO.

## ACKNOWLEDGEMENTS

We would like to thank Maarten H. P. Kole for reading a previous version of the manuscript. This work was supported by a European Research Council Consolidator Grant (ERC CoG 865634) to SH.

## AUTHORS CONTRIBUTION

A.R.J., S.M.S., and S.H. conceptualized the study; A.R.J. and T.T. acquired electrophysiological data; A.R.J. and S.H. analyzed electrophysiological data; M.K. and I.B. performed and analyzed electron microscopic data; J.A., A.R.J., and S.H. devised analysis of loose-seal recorded data; B.B. and S.H. developed the electric circuit model; A.R.J. and S.H. tested amplifier performance by the electric circuit model; A.R.J., J.E., S.M.S., and S.H. interpreted the data and wrote the manuscript. A.R.J. and S.H. generated the figures. All authors approved the final version of this manuscript.

## Supplemental Information

### Supplementary Figures 1–5

**Figure S1.**
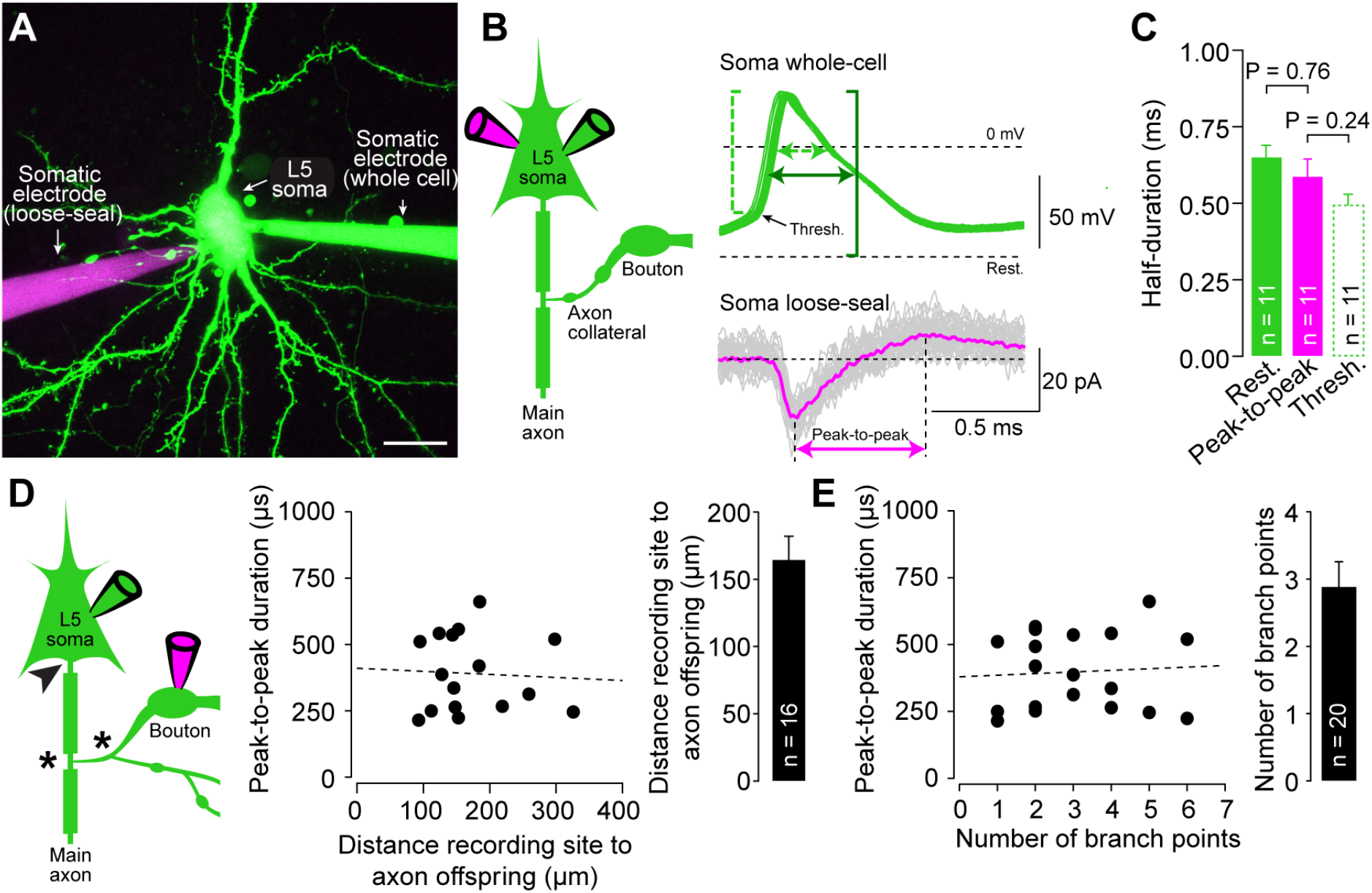
Loose-seal recordings reliably reflect spike duration and indicate stable axonal spike propagation, related to Figure 1. **(A)** 2P-image of a simultaneous somatic whole-cell (green pipette) and loose-seal recording (magenta pipette) from a neocortical layer 5 pyramidal neuron. Scale bar 15 μm, maximum z-projection of a stack of 69 images, z-step size 1 μm. **(B)** *Left:* Pictogram of the recording configuration. *Right:* Overlay of 31 evoked somatic action potentials (green) in whole cell-mode and action potential-evoked somatic currents simultaneously recorded in loose-seal mode (magenta). Somatic action potential amplitude from resting membrane potential (dark green) and action potential threshold (light green, broken line). Action potential half-durations at corresponding half-amplitudes (green arrows) and peak-to-peak duration in somatic loose-seal recording (magenta arrow) indicated. **(C)** Half-duration of somatic action potentials quantified from resting potential and action potential threshold related to peak-to-peak duration of simultaneously recorded currents in simultaneous somatic loose-seal recordings (bar graphs as mean ± SEM, color code as in **b**, n indicates number individual paired somatic recordings). **(D)** *Left:* Pictogram of a paired bouton-soma recording illustrating the offset of the axon proper from the soma (arrowhead) and axonal branch points between axon offset and bouton recording site (asterisks). *Middle:* Peak-to-peak duration of loose-seal recorded currents over distance between bouton recording site and axon offset (broken line as linear fit). *Right:* Distance of bouton recording site from axonal offset (bar graph as mean ± SEM, n indicates number of paired bouton-soma recordings). **(E)** *Left:* Peak-to-peak current duration over number of axonal branch points between bouton recording site and axon offset (broken line as linear fit). *Middle:* Peak-to-peak duration of loose-seal recorded currents over branch point number between bouton recording site and axon offset (broken line as linear fit). *Right:* Branch point number of between bouton recording site from axonal offset (bar graph as mean ± SEM, n indicates number of paired bouton-soma recordings).

**Figure S2.**
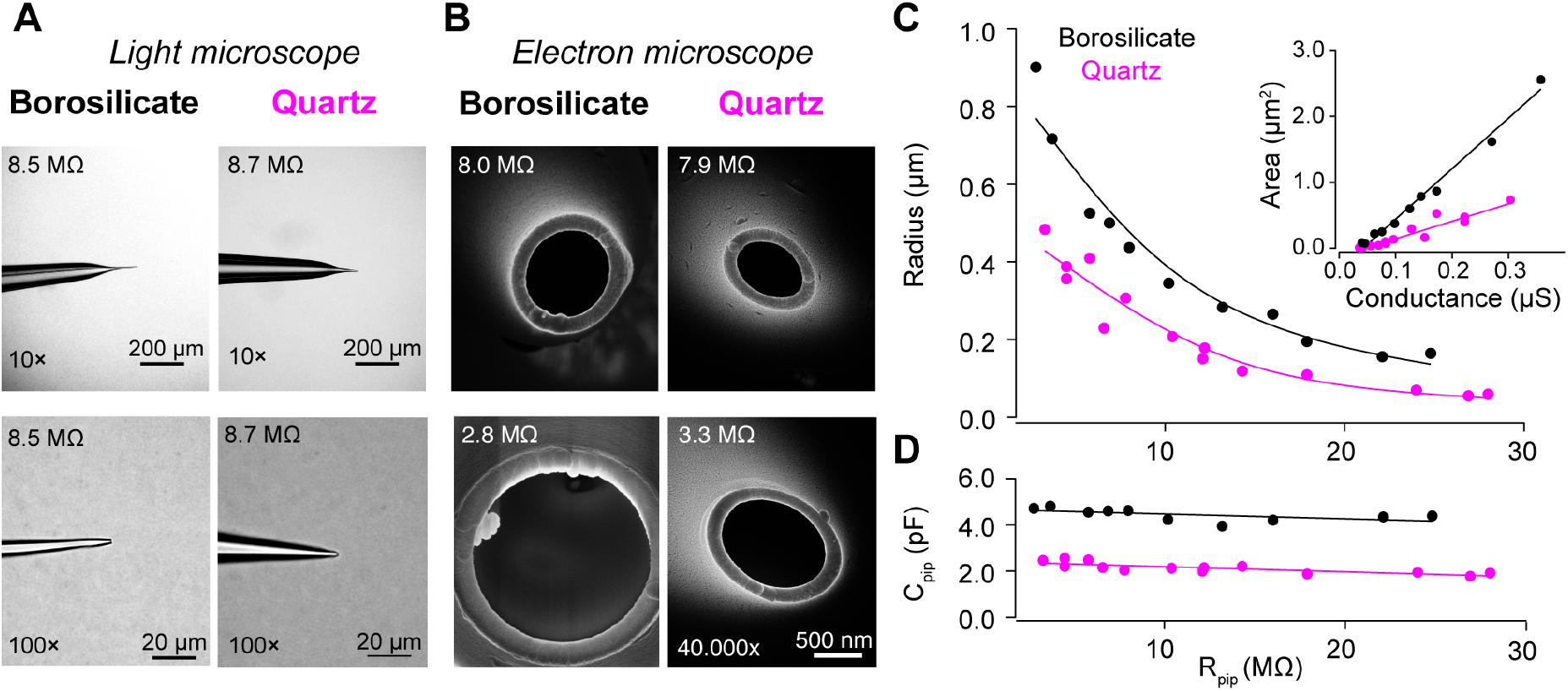
Borosilicate and quartz glass pipettes show different geometrical and electrical properties, related to Figure 2. **(A)** *Left:* Light microscopic images of example borosilicate pipettes at 10x and 100x magnification. *Right:* Light microscopic images of example quartz pipettes at same magnification. **(B)** *Left:* Electron microscopic images of borosilicate pipette tips of various pipette resistances. *Right:* Electron microscopic images of borosilicate pipette tips **(C)** Radius of the tip opening for borosilicate and quartz glass pipettes (in black and magenta, respectively) of different pipette resistances. Solid lines are smoothed spline interpolations to pipette radius. Inset: Pipette tip opening area over pipette conductance with linear fits. Pipette orifice area over conductance was fitted linearly and the radiuses for different pipette resistances were fitted by smooth spline interpolation. **(D)** Pipette capacitance for borosilicate and quartz glass pipettes (in black and magenta, respectively) of different pipette resistances. Solid lines are linear fits to pipette capacitance. Pipette resistances and capacitances were recorded from sister pipettes of those imaged by electron microscopy.

**Figure S3.**
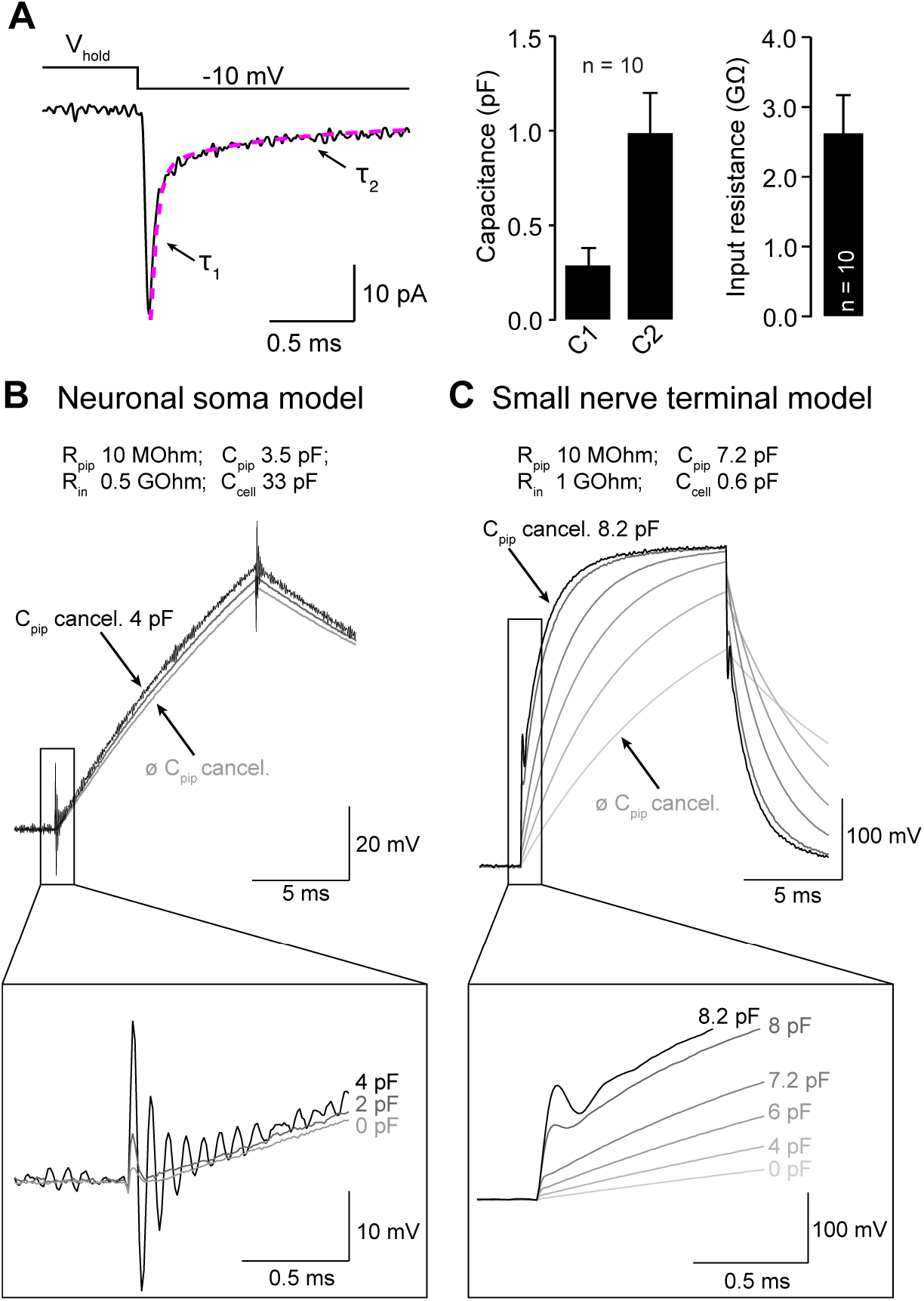
The small capacitance of *en passant* boutons precludes conventional pipette capacitance determination in current-clamp mode, related to Figure 3. **(A)** *Left:* Current obtained from the difference between current responses in cell-attached mode and immediately after establishing whole-cell mode from boutons superimposed with a bi-exponential fit (magenta broken line). *Right:* Derived passive parameters for small boutons. **(B)** *Top:* Increasing pipette capacitance cancellation in current-clamp mode in a neuronal soma model circuit leads to voltages oscillations clearly separable from slower voltage changes of the somatic whole-cell capacitance. *Bottom:* Expansion of the initial voltage change upon current injection for different pipette capacitance (C_pip_) cancellations **(C)** *Top:* Current injection-evoked voltage responses in a small bouton model (same as in Figure 3) for different pipette capacitance cancellations. *Bottom:* Enlarged initial voltage change upon current injections.

**Figure S4.**
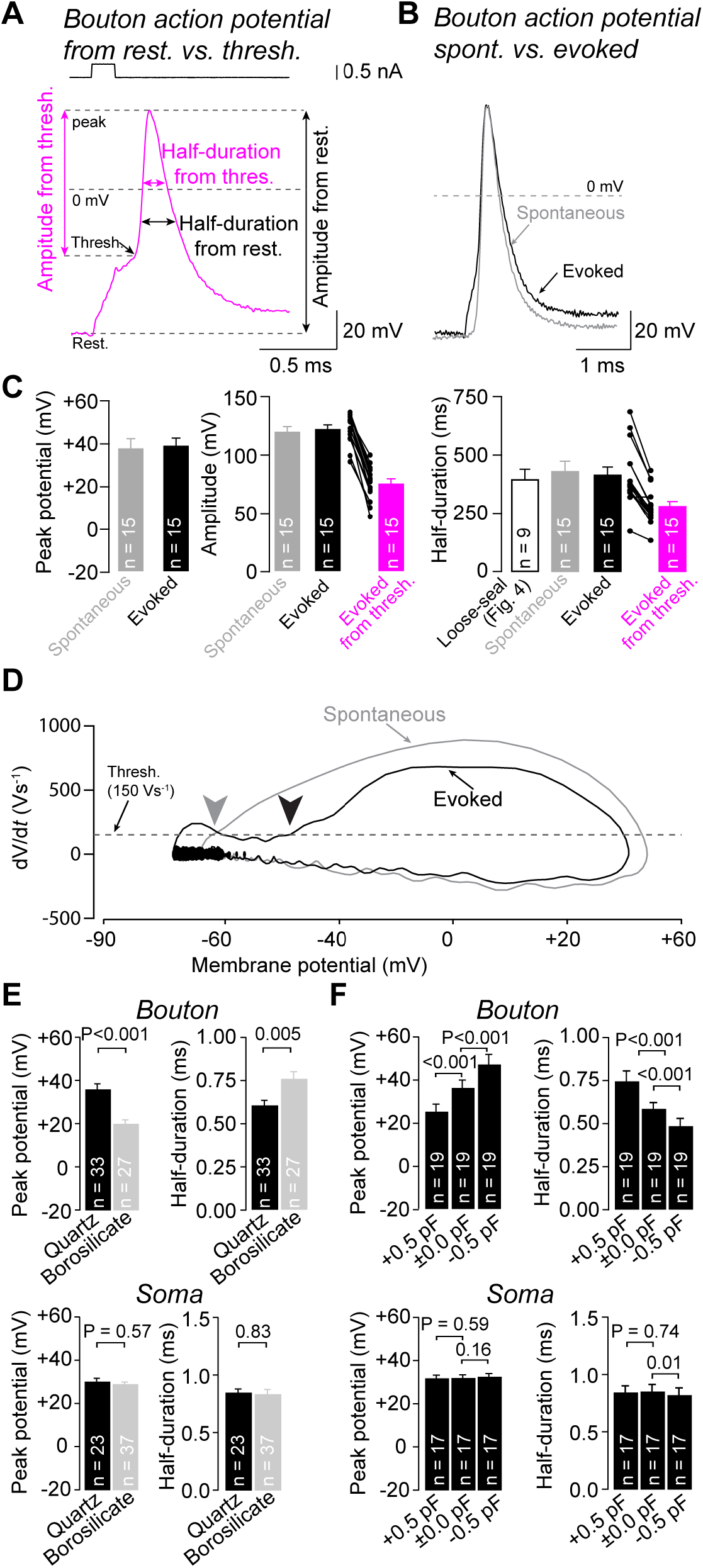
Bouton action potentials are unperturbed by current-injection but strongly affected by absolute pipette capacitance and capacitance compensation, related to Figure 5. **(A)**Illustration of action potential amplitude and half-duration quantified from resting membrane potential (black) and action potential threshold (magenta) of a bouton action potential evoked by current injection with a quartz glass pipette in cultures. **(B)** Overlay of example spontaneous (black) and evoked action potentials (grey) recorded from the same bouton with a quartz glass pipette in cultures. Action potentials were aligned to the action potential peak. **(C)** *Left:* Peak potential of spontaneous and evoked action potentials. *Middle:* Amplitude of spontaneous and evoked action potentials. *Right:* Half-duration of spontaneous and evoked action potentials (bar graphs as mean ± SEM, n indicates the number of individual bouton recordings or paired bouton-soma recordings). **(D)** Phase-plane plot of the evoked and spontaneous action potential shown in **b**. Voltage a which action potential threshold is crossed (defined as 150 Vs^-1^) indicated by arrow heads (color code as in **b**). **(E)** *Top:* Average peak potential and half-duration of action potentials recorded from boutons with quartz (black) or borosilicate glass pipettes (grey). *Bottom:* Average peak potential and half-duration of somatic action potentials recorded with quartz (black) or borosilicate glass pipettes (grey). All recordings with Multiclamp 700A amplifier. **(F)** *Top:* Average peak potential and half-duration of action potentials in boutons when varying capacitance compensation (±0.5 pF). *Bottom:* Average peak potential and half-duration of somatic action potentials when varying capacitance compensation (±0.5 pF).

**Figure S5.**
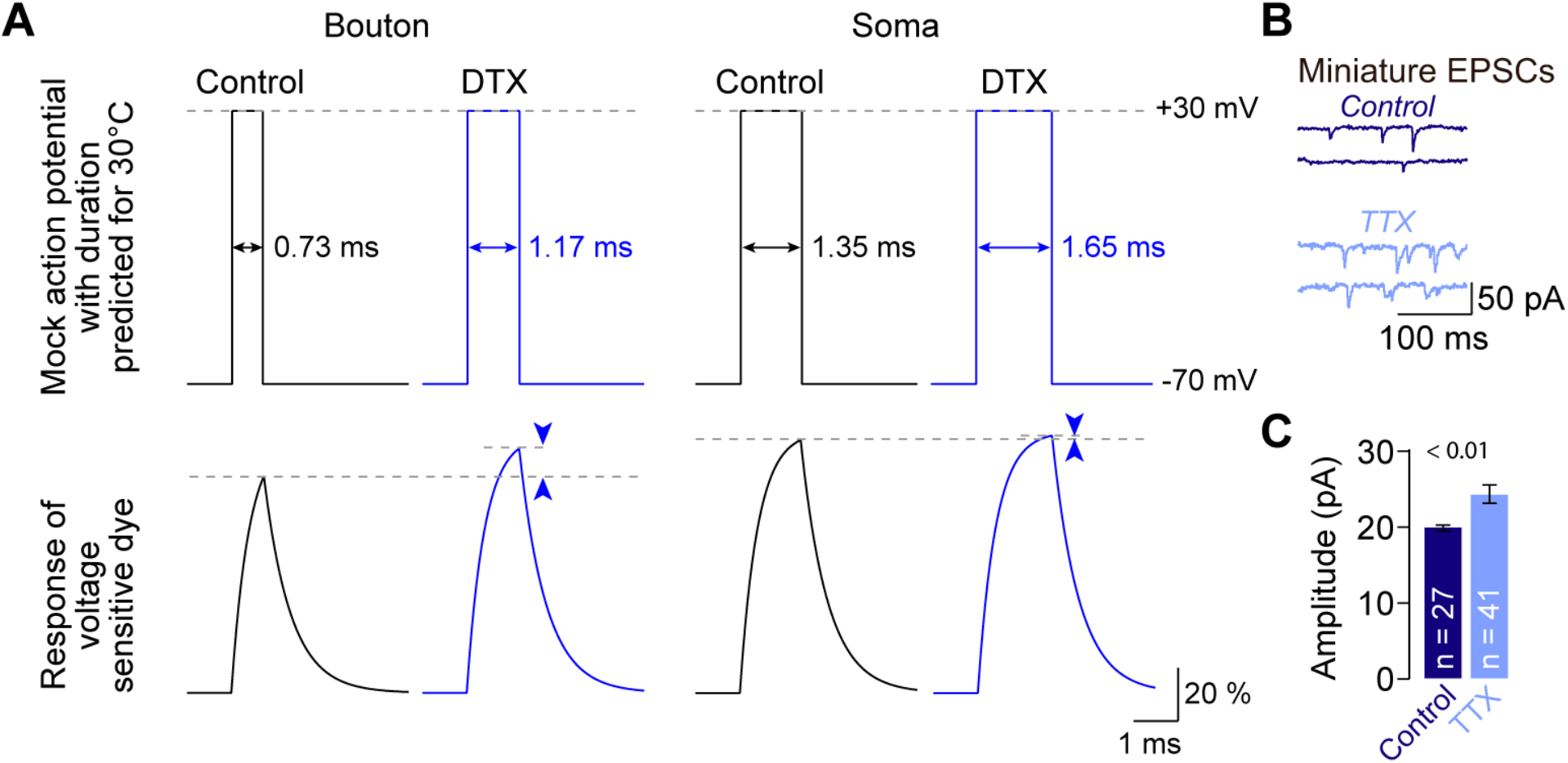
Filtering of spike waveform imposed by voltage sensitive dye kinetics and scaling of postsynaptic miniature events by chronic activity blockade, related to Figure 6. **(A)** Using genetically-encoded voltage sensors, a study (Hoppa et al., 2014) reported small amplitudes of presynaptic action potentials in cultured hippocampal neurons that increases upon pharmacological blockade of Kv channels (recordings at 30°C). Different methods of recording action potentials might account for the difference to action potential reported here. Mock action potentials (*top*) with durations predicted for 30°C from our data measured at 36 °C (assuming a Q_10_ value of 2.2, see *methods*) were used to calculate fluorescence response of the employed voltage sensitive dye (Archaerhodopsin, *bottom*) according to published on- and off-response rates (Bando et al., 2019; Maclaurin et al., 2013) exponential rise and decay time constants were 0.4 and 0.6 ms for steps from −70 to +30 mV and from +30 to −70 mV, respectively). Broadening bouton action potentials (as occurring following potassium channel block; cf. Figure 6) permitted the voltage-sensor enough time to progress further towards steady-state and consequently increased the apparent fluorescence response by ~10 %. In contrast, the slower somatic action potential (Figure 5d) allowed the fluorescent signal greater time to reach steady state and resulted in an apparently larger amplitude. Consequently, action potential broadening by DTX had minimal effect on the somatic fluorescence response or spike amplitude. Small action potential amplitude and the apparent regulation by potassium channels therefore results from the slow kinetics of the employed voltage-sensor that is predicted to differentially distort rapid presynaptic and slower somatic action potential waveforms based on its kinetics (Bando et al., 2019; Maclaurin et al., 2013). **(B)** Example recordings of miniature excitatory postsynaptic currents (mEPSCs) in somatic recordings from cultured neurons under control conditions and after exposure of cultures to TTX (2 μM in culture medium for 48h prior to recordings). **(C)** Mean mEPSC amplitude after exposure to TTX normalized to control conditions (n indicates number of individual somatic recordings)

